# Early Arousal Signals Drive Reward Learning and Subsequent Choice Behaviour

**DOI:** 10.1101/2025.11.25.690494

**Authors:** Kitti Bán, Elsa Fouragnan, Marios G. Philiastides

## Abstract

Learning to reinforce rewarding actions and avoid repeated mistakes is crucial for survival in dynamic environments. Yet, it remains unclear how distinct neural signals coordinate to implement reward-based decision-making and behavioural adjustment. We obtained simultaneous electroencephalography (EEG) and pupillometry during a probabilistic reversal learning task. Leveraging single-trial EEG, we first replicate the presence of two feedback-locked neural representations; an early signal previously linked to alertness and switching behaviours following negative feedback and a late signal associated with value updating and reward learning. Using single-trial pupillometry, we then show that differences in feedback-evoked pupil responses between positive and negative feedback are driven primarily by negative feedback encoding. Jointly examining these EEG and pupillometry signatures, we show that following negative feedback, increased trial-by-trial coupling between the pupil response and the early, but not the late, EEG signal is linked to increased uncertainty and exploration tendency as well as reduced accuracy and evidence accumulation on the next trial. Consistent with previous research implicating the locus-coeruleus-noradrenaline system in uncertainty signalling and network resets, we propose that when internal estimates of contextual uncertainty are high following negative feedback, an early signal, likely regulated by locus coeruleus activity, implements a network reset in reward learning structures of a later learning signal. This interruption may simultaneously increase the neural gain related to the processing of novel information and decrease the influence of existing representations in reward learning structures, in turn improving performance by creating new, more accurate internal representations of the external world.

**Significance Statement:** The current study jointly examines EEG and pupillometry signatures associated with reversal-learning during reward-based learning. It suggests that when internal estimates of contextual uncertainty are high following negative feedback, an early neural signal, likely regulated by locus coeruleus activity, implements a network reset in reward learning structures of a later learning signal. This interruption may simultaneously increase the neural gain related to the processing of novel information and decrease the influence of existing representations in reward learning structures, in turn improving performance by creating new, more accurate internal representations of the external world.

## Introduction

Do we learn differently from better- or worse-than-expected decision outcomes? Converging evidence implicates the dopaminergic system in guiding value-based learning through signalling reward prediction errors (Fouragnan et al., 2015; Gläscher, et al., 2008; O’Doherty et al., 2007; Schultz et al., 1997). However, a complete characterisation of how this learning process is influenced by other neurotransmitter systems, feedback valence, or uncertainty is still lacking.

In addition to the well-established role of cortical and subcortical dopaminergic structures in learning, Fouragnan and colleagues (2015) used EEG-informed functional magnetic resonance imaging (fMRI) to uncover two spatiotemporally distinct, but interacting, value systems encoding feedback valence; an ‘early’ system, activated by negative feedback and engaging arousal-related structures, and a ‘late’ system, consistent with the human reward network, which was modulated by both positive and negative feedback (increased or decreased, respectively).

Crucially, the early system down-regulated the late system following negative feedback, via a thalamo-striatal coupling. The strength of this coupling predicted participants’ switching behaviour and avoidance learning. These results highlight the importance of fast alertness/arousal signals following undesirable outcomes to attenuate value encoding and facilitate behavioural adjustment towards alternative options (Fouragnan et al., 2015; 2017; 2018; Schultz, 2016; Algermissen et al., 2024).

Here, we hypothesise that phasic responses in the locus-coeruleus-noradrenergic (LC-NA) arousal system (Sara & Bouret, 2012) might play a role in regulating the activity of the early system and facilitating its interaction with the late system, by acting as an interrupt signal leading to network resetting (Bouret & Sara, 2005; Dayan & Yu, 2006; de Gee et al., 2017; Filipowicz et al., 2020; Sara, 2009). Indeed, increased phasic pupil dilation and LC-NA activity have been associated with increased levels of uncertainty and surprise (Colizoli et al., 2018; de Gee et al., 2014; Filipowicz et al., 2020; Lavín et al., 2014; Nassar et al., 2012; Preuschoff et al., 2011; Urai et al., 2017; van Slooten et al., 2017; 2018), exploration (Aston-Jones & Cohen, 2005; Jepma & Nieuwenhuis, 2011; Pajkossy et al., 2017), attentional set shifting (Jepma & Nieuwenhuis, 2011; Pajkossy et al., 2017), and boosted behavioural flexibility (Eldar et al., 2013) in reward learning tasks.

In this study, we collected simultaneous EEG-pupillometry data during a probabilistic reversal learning task to investigate how phasic changes in pupil diameter, a proxy for LC-NA activity (Joshi et al., 2016; Joshi & Gold, 2020; Murphy et al., 2014; Reimer et al., 2016), may covary with early and late feedback system activity. We hypothesised that changes in feedback-related pupil responses would be linked with trial-wise changes in the early EEG signal following primarily negative feedback. Moreover, given the interaction between the early and late systems (Fouragnan et al., 2015), we also expected a knock-on effect on the late system, manifesting as a further covariation of the pupil response with the late EEG signal.

Finally, we utilised neurally-informed drift diffusion modelling (DDM; Ratcliff, 1978; Ratcliff & McKoon, 2008; Wiecki et al., 2013) to investigate the contribution of the early and late feedback systems to successive decisions. Although several studies have explored the computational and neurophysiological mechanisms that transform sensory evidence into a choice (Cavanagh et al., 2011; 2014; Chakroun et al., 2023; Forstmann et al., 2010; Frank et al., 2015; Franzen et al., 2020; Krajbich et al., 2010; Mattes et al., 2022; Nunez et al., 2017; Ratcliff et al., 2009; Turner et al., 2015;), little has been done to investigate how feedback-related representations affect subsequent choices. Specifically, we hypothesised that if the early system implements LC-NA-induced network resets in the late system following negative feedback, subsequent decision-making would be decelerated as evidence is accumulated towards a new, competing hypothesis (i.e., reversal in reward contingencies).

## Materials and Methods

### Participants

A total of 73 student participants of unknown sex took part in the experiment. Data of 10 participants were excluded from successive analyses due to poor behavioural performance. This was evaluated by a binomial test statistic with a significance threshold of *p* = 0.01 in order to determine whether participants were selecting the symbols randomly. Our criteria for pupil data exclusion included excessive blinking compromising the pre-processing pipeline (causing unreliable recovery of data points during the interpolation of blinks) and cases where more than 10% of all epoched pupil trials were determined to be outliers (see section on pupil data analysis below). We also excluded EEG data with excessive noise that led to poor feedback valence (positive versus negative) discrimination. Consequently, we excluded 9 participants based on meeting pupil exclusion criteria, a further 3 participants as a result of inadequate EEG data, and 3 participants that had both poor quality pupil and EEG data. Following the exclusion of 25 participants based on the above criteria, we retained data from 48 participants in our analyses. All participants were right-handed, had corrected-to-normal vision, had no existing psychiatric conditions, and were not taking psychoactive medication at the time of the study. Each participant gave written, informed consent in accordance with the School of Psychology and Neuroscience Ethics Committee at the University of Glasgow.

### Stimuli display

The experiment was carried out through Presentation software (Neurobehavioral Systems, Inc., Berkeley, CA). Crucially, we designed all stimuli to be equiluminant across all trials and trial events of the experiment to ensure that pupil responses were not confounded by changes in luminance (Mathot & Vilotijevic, 2022). Stimulus symbols entailed a line crossing a circle, where the angle of the lines determined stimulus type (delay symbol, and two stimulus symbols; Fig.1a). The symbols were placed on the left and right side of a small vertical line enclosed from the top and the bottom by short horizontal lines. During the feedback screen of each trial, this central figure, surrounded by the delay symbols on each side, was transformed into an arrow pointing upwards or downwards, informing participants of the feedback associated with each trial. A representation of a typical trial in the experiment is displayed on Fig.1a. To reduce pupil data distortions, participants were seated directly opposite the eye-tracker camera. Participants sat in a sound-proof booth with intermediate levels of ambient lighting in order to minimise distractions and discomfort during the task.

**Figure 1.**
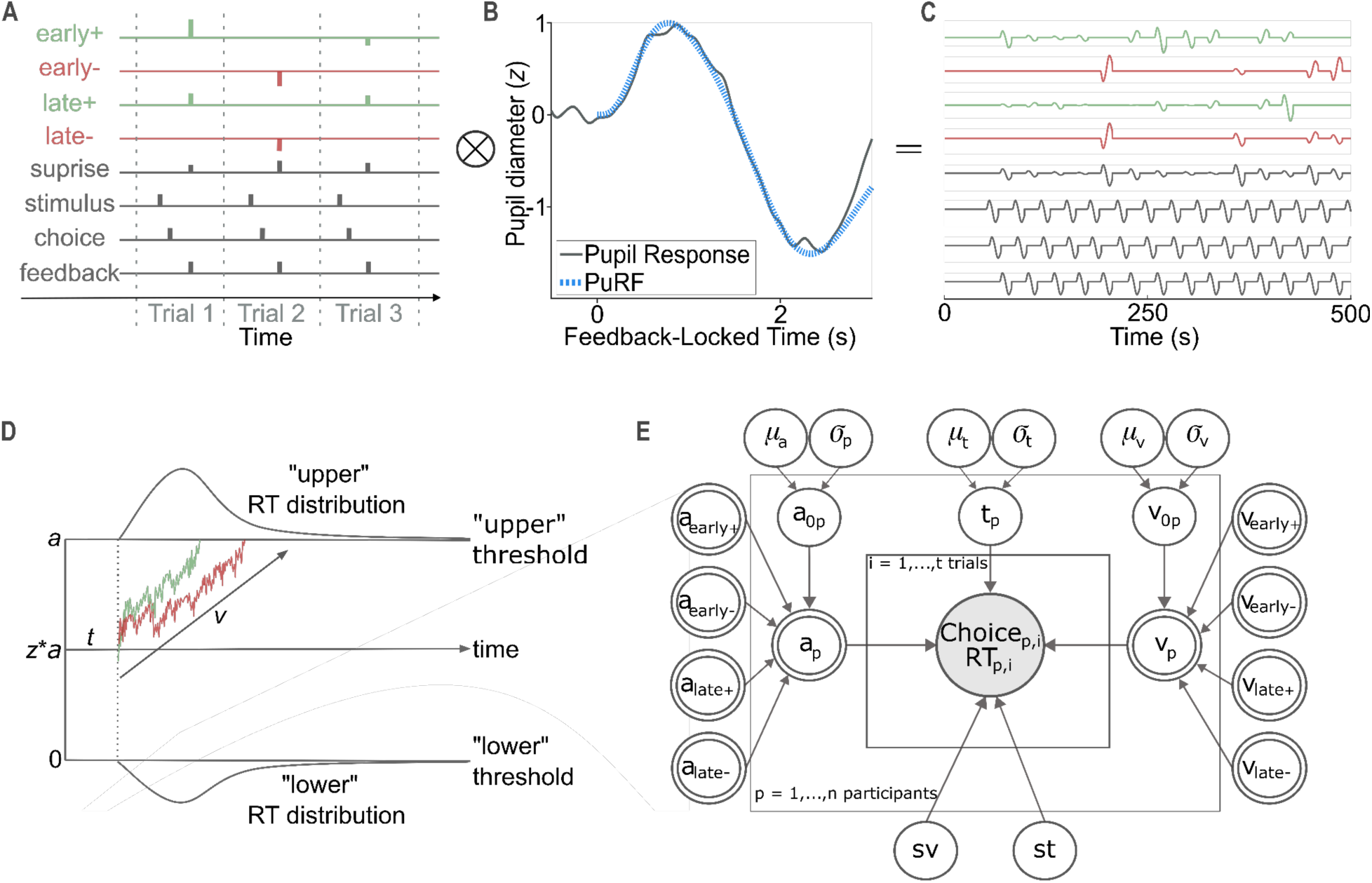
EEG-informed pupil analysis and drift diffusion model. **A.** Our participant-wise GLM predicted the full course pupil data and included 8 regressors. Four parametric regressors were modulated by the amplitudes of the *y_i_* (Eq.1) values derived from our feedback-locked linear discriminant analysis (yielding early negative, early positive, late negative and late positive predictors). To represent the effect of surprise, we created a parametric regressor modulated by the absolute value of the prediction error estimated by our reinforcement learning model. To account for variance linked to other task-related processes, we included non-parametric, boxcar regressors at the times of stimulus display, response, and feedback presentation, the value of which was set to 1 at the appropriate time points. Finally, we convolved each regressor with the **B.** participant-specific pupil response function (PuRF) that was adjusted to each participant’s mean feedback-locked pupil response. **C.** The transposed design matrix (*X^T^*) is shown for the first 500 ms of the experiment, after the predictor indicator functions have been convolved with the participant-specific PuRF. **D.** Hypothesised drift rate effects. Drift trajectories are shown following positive and negative feedback within the drift diffusion modelling (DDM) framework. Evidence is stochastically accumulated over time (*x*-axis) with average drift rate *v* until one of the two boundaries (at 0 and *a*, representing the “upper” and “lower” thresholds, respectively) is crossed, and a response is initiated. The diffusion process begins at the starting point between the two thresholds (marked by the proportion of *a* by *z*). We hypothesised that evidence accumulation (i.e., drift rate) following negative (blue line) compared to positive (red line) feedback will be reduced. Total response time is the sum of the non-decision time *t*, denoting the time taken for stimulus encoding and motor processing, and the duration of the diffusion process. The upper and lower plots represent RT distributions for drift diffusion processes hitting the upper and lower boundaries, respectively. **E.** Our hierarchical drift diffusion model (DDM) with trial-wise neural regressors is shown. Round nodes illustrate continuous random variables and double-bordered nodes illustrate deterministic variables, defined in terms of other variables. Shaded nodes represent observed data (RT, choice), whilst unshaded notes represent latent variables. Participant-specific parameters *t_p_*, *v_p_*, *a_p_*, and *z_p_* are estimated from individuals drawn from a group distribution with inferred mean μ and variance σ. Trial-to-trial variations in the decision threshold *a* and drift rate *v* are determined by the amplitudes of four EEG discriminating components (*y_i_*, Eq.1). Multiple random variables share the same parents and children (e.g., each participant specific threshold parameter *a_p_* shares the same parents that determine the group distribution). The inner plate is over trials *i* and the outer plate is over participants *p*. Trial-by-trial neural regression coefficients were estimated on the group-level only to increase estimation reliability and avoid parameter explosion.

### Reversal learning task

The experiment involved a total of 300 trials, broken down into four blocks and separated by breaks. In each trial, participants had to choose from two stimulus symbols, with the aim to discover the one with the higher reward probability. Participants were informed that changes in reward contingencies may occur during the blocks and that they would need to modify their choice behaviour accordingly. Participants received a fixed compensation of £6 per experimental session as well as a further performance-based payment of up to a maximum of £6.

Each trial commenced with the same delay symbol appearing on both the left and right side of the screen for a random interval between 2 to 3 seconds. Next, two stimulus symbols replaced the delay symbols for 1.25 seconds, during which participants picked one of the symbols by pressing designated buttons on a response box. After signalling their choice, participants saw a second delay screen for 1.5-2 seconds. Choice feedback was issued by an arrow in the middle of the screen signalling either a positive or a negative feedback (i.e., the arrow pointing upwards or downwards, respectively). Fig.1a outlines this sequence of trial events. In case participants failed to choose a symbol on time during stimulus representation, the feedback screen displayed a message reminding participants to make a faster choice next time. Such lost trials were excluded from all analyses.

At any point during the experiment, one of the stimulus symbols was corresponding to a high reward probability of 70%, whilst the other symbol had a reward probability of 30%. Importantly, feedback for each symbol was generated independently (i.e., it was drawn from separated distributions). Participants were naïve about the reward probabilities allocated to stimulus symbols and were instructed to discover the symbol with the higher reward probability through trial and error by considering the feedback in each trial. We determined reversals points in a way that ensured that participants have a sufficient amount of time to learn and exploit the ongoing reward contingencies based on data from previous reversals learning experiments from the lab (Carvalheiro & Philiastides, 2023; Fouragnan et al. 2015, Queirazza & Philiastides, 2024; Westwood & Philiastides, 2024). During the task, participants experienced an average of 9.96 reversals (*SD* = 0.20) with a mean number of 21.37 trials (SD = 5.51, *min* = 15, *max* = 30) between reversals. In order to become familiar with the task, participants practised a block of 75 trials prior to the experiment.

### Reinforcement learning model

We used a model-free reinforcement learning model optimised for reversal learning paradigms (Krugel et al., 2009) to derive trial-by-trial reward prediction errors. In reinforcement learning models, each choice option is assigned an expected reward value *q_A_(i)*, which in turn is used to obtain choice probabilities *p_A_(ti)*. This is derived from choosing option *A* in trial *i*, following the softmax choice rule:

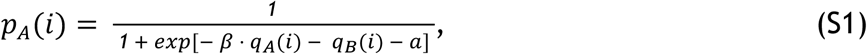

where *ꞵ* is the sensitivity parameter controlling the weight of reward expectations on choice probabilities, *q_A_(i)* and *q_B_(i)* are the expected values of each of the two choice options, and *a* is the indecision point, representing equiprobable choice between the stimuli (Hampton et al., 2006). After the participant selected one option, the observed feedback *r_A_(i)* is compared with the expected reward *q_A_(i)*, where the discrepancy produces the reward prediction error *δ* (RPE):

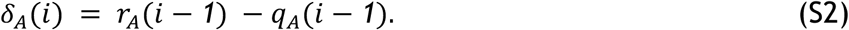

Reinforcement learning models postulate that the deviations expressed by RPEs drive learning as expected choice values are updated as follows:

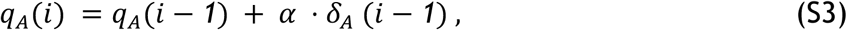

where *α* is the learning rate that controls the influence of the RPE on the updating of the stimulus expected value. The expected value of the unselected stimulus on trial *i* is not updated. A dynamic learning rate *α(i),* instead of a constant learning rate *α*, was used to capture the participant- and trial-wise fluctuations in the task introduced by reversals. This dynamic learning allows for both rapid adaptations after reversal and the stabilisation of behaviour once the better option has been discovered. It is modulated by the slope *m* of the smoothed and unsigned RPE according to the following update rule:

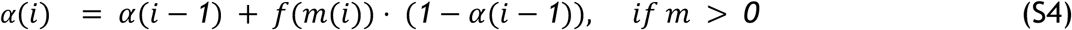

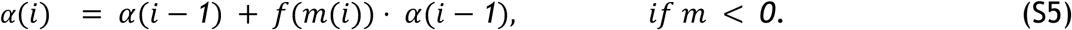

Accordingly, the learning rate increases when the slope of the unsigned RPE is positive and decreases when the slope is negative. The slope was estimated over the smoothed and unsigned RPEs, where smoothing is determined as follows:

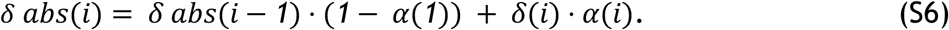

Here, *α*(1) serves as the learning rate for the first trial but also represents the learning rate used to update the unsigned RPE. Consequently, high values for *α*(1) suggest that only the most recent RPEs determine present estimate of the RPE weight. The PE slope was normalised over the last two RPEs in order to produce a slope that is independent of the scale of payoffs:

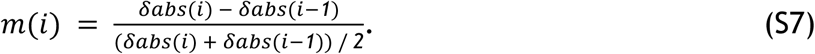

Finally, the weight of the slope of the RPE on the learning rate was transformed by a double sigmoid function that allows for the slope to take on values between 0 and 1, according to

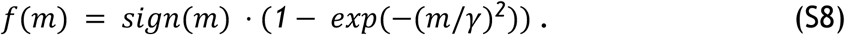

Here, the free parameter *γ* controls the extent to which the RPE slope influences the learning rate. Therefore, if *γ* takes on a high value (*γ* > 3), the updating of the dynamic learning rate is negligible, so that the learning rate becomes a constant that is guided by *α*(1).

For each participant, we estimated four parameters: the sensitivity parameter of choice *ꞵ*, the dynamic learning rate *α*(i), the sensitivity of the learning rate with regards to the RPE slope *γ*, and the indecision point *a*. We used the following predetermined starting points for the above parameters:

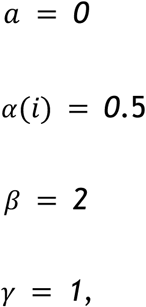

These values constituted the starting points of the maximum likelihood estimation fitting procedure for sets of parameters *θ_j_*, such that

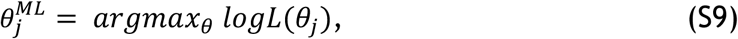

where the likelihood *logL* was calculated according to

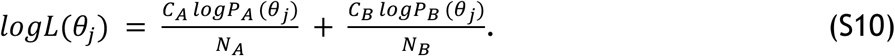

Here, *P(θ_j_)* stands for the choice log probability given the model parameters *θ_j_, C* is a binary vector for observed choice, *N* is the number of observed choices, and the subscripts mark each of the available two choices. During the optimisation procedure, we constrained the free parameters according to:

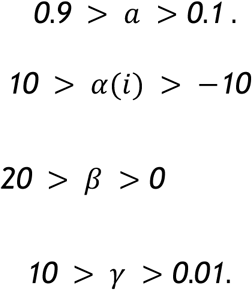

Lastly, to obtain estimates of trial-by-trial estimates of RPEs and dynamic learning rates, the final set of parameters were reintroduced into the reinforcement learning algorithm.

The value of the free parameter *γ* was estimated as < 3 for the majority of participants (35 out of 48), indicating that learning was regulated by a dynamic learning rate. Accordingly, in subsequent analyses, we used RPE estimates derived by the algorithm utilising dynamic learning rates. To illustrate the model’s goodness of fit, we binned the participant-specific choice probabilities into ten groups based on distribution quintiles and calculated the participant-specific average choice probability for each bin. This allowed us to recover the participant- and group-wise mean observed choice probabilities as well as the corresponding Pearson’s correlation coefficient (*r* = 1, *p* = 0), both of which are shown on Fig.1B.

### EEG data acquisition and pre-processing

We simultaneously acquired continuous EEG and pupillometry data from participants performing the experimental task in an electrostatically shielded and sound-attenuated booth. We used a 64-channel EEG amplifier system (BrainAmps MR-Plus, Brain Products GmbH, Germany) with Ag/AgCl scalp electrodes situated following the international 10-20 system on an EasyCap (Brain Products GmbH, Germany). A chin electrode served as a ground and all EEG channels were referenced to the left mastoid. Input impedance was adjusted to under 20 kΩ. Data were recorded in Brain Vision Recorder (BVR; Version 1.10, Brain Products, Germany) at a sampling rate of 1000 Hz and subjected to online (hardware) filtering by an analog band-pass filter of 0.016 - 250 Hz. Experimental event trigger codes of participant responses, stimulus and feedback presentation were synchronised with the EEG data and collected via Brain Vision Recorder. These data were stored for offline analysis in MATLAB (version 2018b, The Mathworks Inc., 2018). We implemented a band-pass filter with cutoff frequencies between 0.5 and 40 Hz to the data to eliminate slow direct current drifts and high frequency noise. Beyond this, a notch filter at 50 Hz was used for line noise reduction. Finally, the data were re-referenced to the average of all electrodes.

To remove eye-movement artefacts, participants were asked to complete an eye-movement calibration task before the main experiment. During this task, participants were required to blink repeatedly while viewing a fixation cross at the centre of the screen, after which they were asked to make a number of horizontal and vertical saccades in accordance with the location of the fixation cross. The timing of these visual cues was recorded, allowing us to identify linear EEG sensor weights linked to blinks and saccades using principal component analysis (Parra et al., 2005), which in turn were projected onto and subtracted out from the broadband data from the main task. Due to the lack of eye calibration data for two participants, we employed independent component analysis implemented in EEGLab (Delorme et al., 2007), which utilised the data from the main task to remove eye-movement artefacts.

### Multivariate EEG data analysis

We aimed to exploit the single-trial variability in the EEG-derived, temporally-specific representations of feedback valence to create parametrically-modulated regressors for our EEG-informed behavioural and pupil analyses, designed to disentangle brain networks associated with reward learning. Similar to previous work (Fouragnan et al., 2015; 2017; Franzen et al., 2020; Parra et al., 2005, Sajda et al., 2009), we used a linear multivariate classifier to EEG data locked to the time of feedback, using a sliding window method. This allowed us to identify a projection (*y_i_*) of the multidimensional EEG signal, *x_i_(t)*, where i = (1…N trials), within a short time window that achieved the greatest level of discrimination between positive and negative feedback trials. All time windows had a width of 60 ms and the window centre was moved from −100 ms to 600 ms relative to feedback onset, in increments of 10 ms. Thus, our discriminator was trained to map positive component amplitudes to positive feedback and negative component amplitudes to negative feedback.

Specifically, the classifier produced a 64-channel spatial weighting *w(τ)* via logistic regression (Parra et al., 2005) that maximally discriminated between positive and negative feedback trials within each time window, arriving at the one-dimensional projection *y_i_(τ)*, for each trial *i* and time window *τ*:

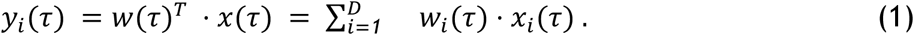

Here, *T* is the transpose operator, *D* is the number of EEG channels, *y_i_(τ)* is a vector of single trial discriminator amplitudes (1 x trials), *w(τ)* is a weight vector with a length corresponding to the number of EEG channels (1 X 64), and *x_i_(τ)* is a data matrix (Channels X Trials/Samples).

We assessed the performance of the discriminator for each time window using the area under a receiver operating characteristic curve, specified as the *A_z_* value, coupled with a leave-one-trial-out cross-validation method to control for overfitting (Philiastides & Sajda, 2006; Philiastides, Ratcliff, & Sajda, 2006). Accordingly, we used *N*-1 trials for each iteration to determine a spatial filter *w*, which was in turn applied to the left-out trial to derive out-of-sample discriminant component amplitudes, *y_i_*, and calculate the *A_z_* curve. To evaluate the significance of the discriminator, we applied a bootstrapping technique that implemented leave-one-out tests following a randomisation of positive and negative feedback trial labels. Repeating this randomisation procedure a thousand times (at 200 ms following feedback presentation to ensure reliable discrimination performance) allowed us to derive a random probability distribution for *A_z_* values, which we in turn utilised as the reference to estimate the original *A_z_* value leading to a significance level of *p* < 0.01 on the random distribution (population *A_z_sig* = 0.598). All the above discriminant analysis steps were performed on the participant level, whereby each participant served as their own replication unit.

The linearity of this model allowed us to derive the scalp topographies of the discriminating components resulting from Eq.(1) by estimating a forward model according to:

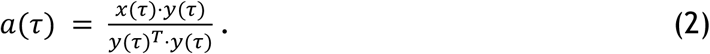

Here, for each time window τ, the discriminating component y(τ) is vectorised (1 X Trials) and the EEG data *x(τ)* is shown in a matrix form (Channels X Trials). The forward model *a(τ)* can be considered a scalp plot and interpreted as the coupling between the discriminating component amplitudes and the observed EEG. Thus, *a(τ)* represents the electrical coupling of the discriminating component *y(τ)* that describes the majority of activity in *x(τ)*. Consequently, a stronger coupling is associated with a low attenuation of the component and reflects the intensity of *a(τ)*. Using this approach, we merged information spatially across the multidimensional electrode space, allowing us to retain the trial-by-trial variability in the discriminating component and increase the signal-to-noise ratio of our data.

In order to establish the early and late valence components, we identified peak discrimination performance in pre-defined time windows for each component (150-290 ms and 300-550 ms for the early and late component, respectively) based on the results reported by Fouragnan and colleagues (2015), while ensuring the participant-specific topographies for each component consistently matched those in previous results. Specifically, to evaluate the spatial characteristics of the discriminating activity, we examined the resulting scalp plots, using the corresponding participant-specific *a(τ*) vectors. In subsequent analyses, we utilised the participant-specific discrimination amplitudes (*y_i_*) linked to the individually-selected early and late components within each trial as an index of how decision feedback is encoded in individual trials (Fouragnan et al., 2015; 2017; Franzen et al., 2020).

### Pupil data acquisition and pre-processing

We used the Tobii Pro x3-120 system (Tobii AB, Stockholm, Sweden) to record participants’ pupil responses from both eyes at 40 Hz during the experiment. Pupil data were pre-processed and analysed in MATLAB (version 2018b, The Mathworks Inc., 2018). We used the averaged pupil diameter across both eyes during all our analyses. We removed invalid data points, resulting from cases when the eye-tracker was unsuccessful in detecting the eyes. Lost data resulting from eye blinks was replaced by linearly interpolated values in the range of −100 to +100 milliseconds around missed events. Finally, we applied a bandpass filter of 0.01 - 4 Hz to remove measurement noise resulting from non-physiological sources (Hoeks & Levelt, 1993; van Slooten et al., 2017) and *z*-scored each participant’s data.

### Analysis of the feedback-evoked pupil response

To derive the participant-wise feedback-evoked pupil response, we epoched the pupil data with a baseline correction of 500 ms relative to feedback presentation and considered data up until 3000 ms following feedback display. We computed the average pupil response for each 25 ms interval within each epoch, yielding 141 data points per trial. To account for outliers, we removed trials in which the pupil response was more than 3 standard deviations away from the mean pupil response or the trial standard deviation fell below the 0.11 cutoff value (the latter was performed to exclude trials with missing data). Using this outlier-removal method, on average, 7.65 trials *(SD* = 6.32) were removed from each participant’s dataset (2.55% of all trials).

To assess the difference in the evoked pupil response due to feedback valence, we calculated the population mean evoked response separately for positive and negative feedback trials for each 25 ms interval within our epoched time frame. All trials were individually baseline-corrected by calculating the mean pupil response in the 500 ms time window preceding feedback presentation and then subtracting this baseline value from at each point in the pupil time series. Subsequently, we compared the mean response for positive and negative trials in each of the intervals using non-parametric cluster-based permutation *t*-tests (Maris & Oostenveld, 2007) for dependent samples. The permutation tests were conducted via *permutest* in MATLAB (Gerber, 2025) and implemented a thousand permutations against a 0.05 significance level.

### EEG-informed pupil analysis

Next, we probed the hypothesised link between pupil dilation, a proxy for LC-NA activity (Joshi et al., 2016; Joshi & Gold, 2020; Murphy et al., 2014; Reimer et al., 2016), and the early system. As the pupil response at any given point may reflect the influence of various ongoing internal signals linked to perceptual and cognitive processing, merely contrasting the feedback-evoked pupil response between positive and negative feedback cannot disentangle the differences associated with specific cognitive processes. In order to link the pupil response to distinct internal signals evoked by specific events, we used the general linear model (GLM) approach (de Gee et al., 2014; Denison et al., 2020) to model the linear combination of the early and late component-associated pupil responses. In this model, the pupil responses linked to components are the internal signal time series linked to individual trial events.

Specifically, we created single-trial predictors reflecting the participant-specific discriminating amplitudes (*y_i_*, Eq.1) of the early and the late components for each trial, both of which were further broken down into positive and negative feedback. We used the resulting four regressors, i.e., early positive, early negative, late positive, late negative, to explain the unique contribution of these two systems in explaining the feedback-related pupil response. Therefore, our regression analyses effectively removed overall unspecific valence effects across positive and negative feedback (Westwood & Philiastides, 2024), and only the trial-by-trial variability within each positive and negative component was used to predict behaviour. This allowed us to test the hypothesis that early system activity is associated with the LC-NA system, and thereby contributing to the increased feedback-related pupillary effect following negative feedback.

To remove nonspecific arousal effects across positive and negative feedbacks, we created a parametric salience predictor, the amplitudes of which were modulated by the unsigned reward prediction error (RPE) obtained from our reinforcement learning model (Eq.S2). To absorb any nonspecific effects related to the presentation of the feedback, stimulus onset, and the time of the decision, we created boxcar predictors, with amplitudes set to 1, at the time of feedback, stimulus display, and choice, respectively (van Slooten et al., 2018). We used the root mean square error (RMSE) to determine whether a more complex model with all the above predictors better explained the pupil data than a simpler model with only the four EEG predictors.

All of the above regressors were convolved with participant-specific pupil response functions (PuRF), which were modelled based on the canonical Erlang gamma function (Denison et al., 2020; Hoeks & Levelt, 1993). The PuRF spanned from −500 to 3000 ms relative to trial events. The width, peak time, and undershoot of each PuRF was adjusted to match the mean participant-wise feedback-evoked pupil response (Fig.2b). Furthermore, the PuRF for each participant was normalised to a maximum value of 1, so that if a regressor component amplitude value of 1 in each predictor amounted to a 1% increase in pupil size compared to baseline (Denison et al., 2020). Using these 8 predictors and the pupil time series as the dependent variable, we fit a linear model to each participant’s data using MATLAB’s *robustfit* function. We employed *t*-tests to evaluate the significance of the effect of the 4 EEG-derived components on the pupil response. As such, for each regressor type (earlyPos, earlyNeg, latePos, lateNeg), we utilised a one-tailed *t*-test to assess whether the group of participant-wise coefficients came from a distribution with mean 0.

**Figure 2.**
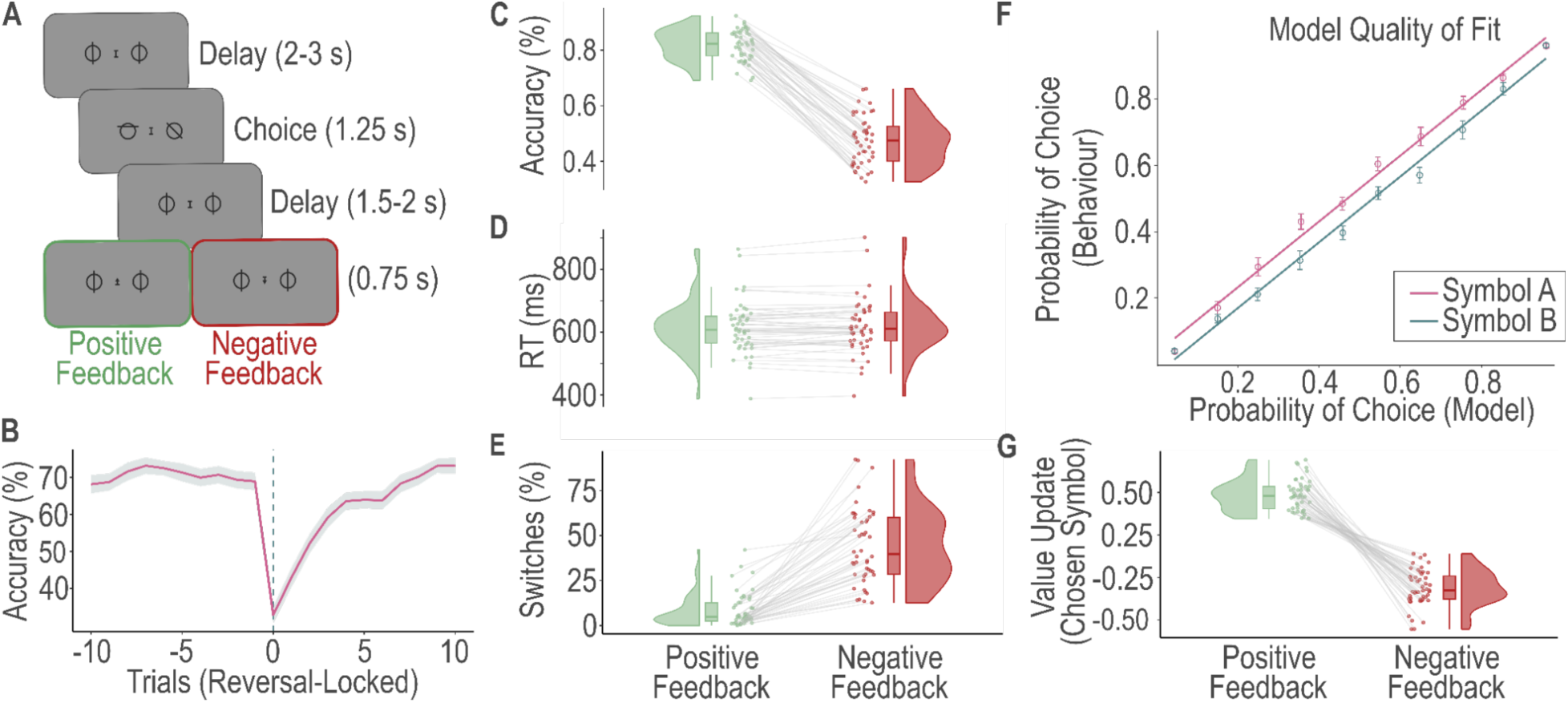
Experimental task, model-free and model-based behaviour. **A.** Representation of a typical trial from the experiment. Each trial commenced with a random delay of 2-3 seconds. Next, participants had 1.25 seconds to choose from two abstract symbols, with the goal of selecting the one with the higher reward probability (70% versus 30%). Following a further random delay of 1.5-2 seconds, an arrow in the middle of the screen indicated either a positive or a negative (i.e., an arrow pointing upwards or downwards, respectively) feedback for 0.75 seconds. Participants performed 4 blocks of 75 trials each. **B.** Population accuracy as a function of trial position locked to reversal points. The shaded error bars show across-participant standard errors. **C.** Population accuracy, defined as selecting the symbol with the higher reward probability, broken down into trial type (positive or negative feedback). **D.** Population reaction times broken down into trial type (positive or negative feedback). **E.** Population probability of switching symbols on the next trial following positive and negative feedback. **F.** Illustration of the behavioural model fit. Predicted choice-probabilities from the reward learning algorithm (*x*-axis), which utilised a softmax method (binned into ten bins, with a bin size of 0.1, and averaged across all participants for both symbols A and B), strongly corresponded to observed choices (*y* axis), derived as the proportion of trials in which participants chose symbol A or B. For each bin, error bars represent the standard error of the mean around the observed choice probabilities. The solid lines show the degree of correlation between the observed and predicted choice probabilities for each of the choice options. **G.** Value updating from trial(*t*) to trial(*t*+1) derived from the reinforcement learning model (Eq.S3), broken down by trial type (positive and negative feedback trials).

**Figure 3.**
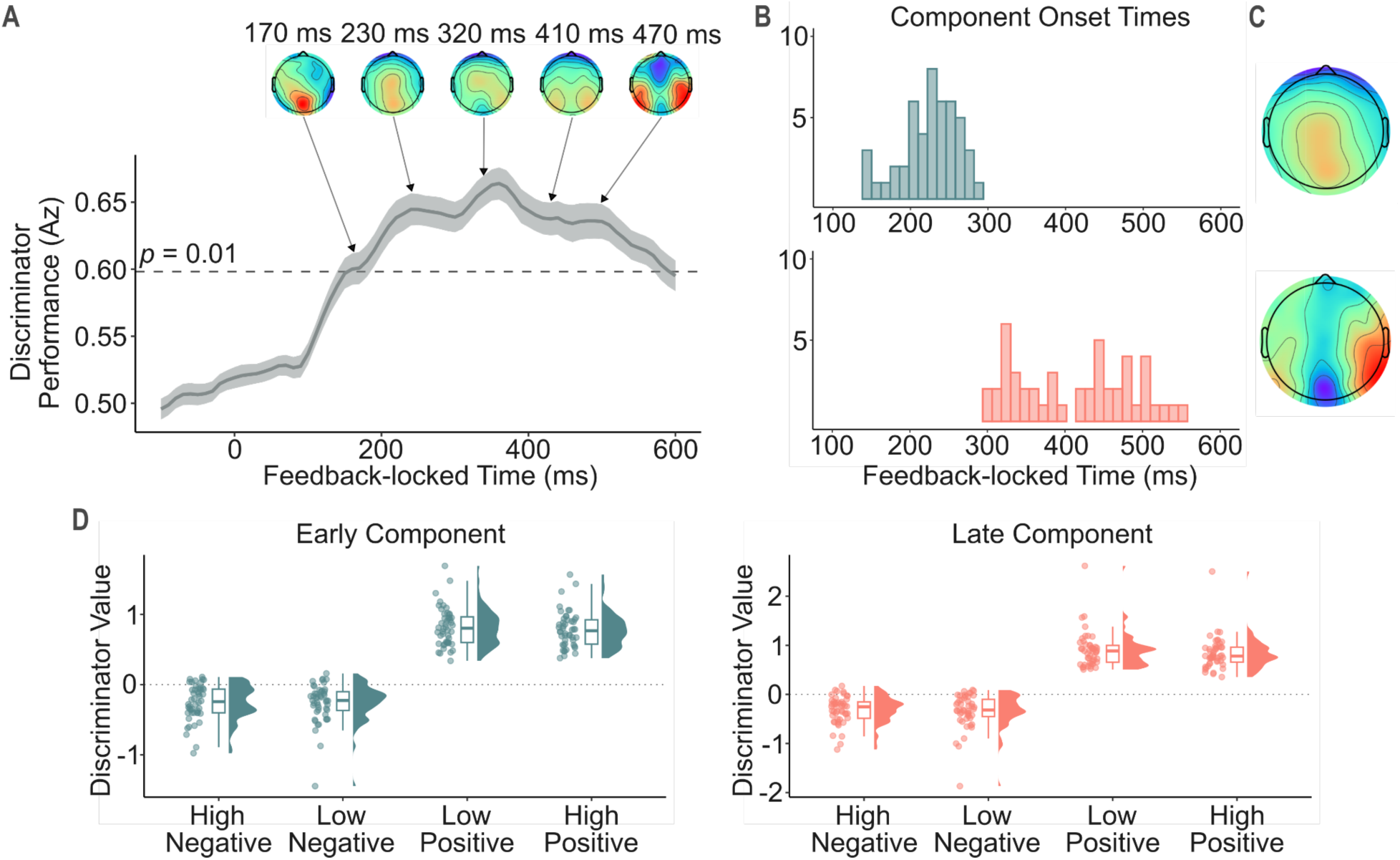
Single-trial EEG analyses. **A.** Single-trial discriminator performance (cross-validated *A_z_*) during valence discrimination (positive vs negative feedback) of the feedback-locked EEG signal, averaged across participants (*n* = 48). The horizontal line illustrates the *A_z_* associated with a significance level of *p* = 0.001, approximated by a bootstrap test. We focused on recreating the early and late valence components that were established by Fouragnan and colleagues (2015). The shaded error band shows across-participant standard errors. The scalp maps on top represent the evolution of the population feedback-related EEG response at significant peak points of the population *A_z_* curve. **B.** Illustration of participant-specific early (top) and late (bottom) component onset times. **C.** Scalp maps illustrating the across-participant spatial topographies of the early (top) and the late components (bottom). **D.** Mean participant-specific discriminator output values (y_i_, Eq.1) linked to the early (left) and late (right) components are binned by RPE magnitude, derived from our reinforcement learning model. Both components revealed predominantly categorical response characteristics, without modulation by RPE magnitude.

To further confirm the GLM results obtained above, we implemented a linear mixed effects model utilising MATLAB’s *fitlme* function. In this model, we treated participants as random effects by specifying participant-specific model intercepts as well as slopes for the four EEG-derived components (Baayen et al., 2008). Similar to the participant-specific GLMs, we used the above 8 regressors to predict the full-course pupil data. To evaluate the degree to which each regressor contributed to the pupil response, we utilised the built-in coefficient evaluation function in *fitlme*, which determined the *p-*values associated with the *t*-statistic for two-sided hypothesis tests.

Next, we aimed to determine how the component regression coefficients with significant predictive power related to behavioural markers, including choice ambiguity, exploration tendency, and mean accuracy. The former was derived from our reinforcement learning model by calculating the absolute difference in probabilities linked to choosing either symbols A (*p_A_*) or B (*p_B_*). As lower values indicate less substantial differences between the likelihood of choosing one option over the other, they index decisions that are more ambiguous (Bland & Schaefer, 2012).

To test the hypothesis that early-system-mediated network resets in the late system increase exploration tendency, we tested whether increased coupling between the early system and the pupil response is associated with a boosted exploration tendency. To quantify exploration tendency, we used two different measures. First, we calculated the proportion of decisions in which participants chose the symbol with the lower subjective value (Daw et al., 2006; Harada, 2020; Warren et al., 2017), which estimates were derived from our reinforcement learning model. As an additional measure, we utilised the inverse temperature parameter obtained from our reinforcement learning model. This free parameter marks exploration-exploitation tendency — the degree to which participants’ decisions are guided by the difference in reward values. Higher values indicate a more substantial influence of symbol value deviations on choice, and therefore mark reduced exploratory propensity (van Slooten et al., 2019).

We used robust bend correlations to measure the association between the four behavioural markers and the regressions coefficients obtained from the EEG-informed pupil GLMs. For this, we utilised the *‘bendcorr’* function in MATLAB from the robust correlation toolbox devised by Pernet and colleagues (2013), which down-weighs bivariate outliers by 20% of all data points in each dimension (i.e., bending constant = 0.2). This function outputs a correlation coefficient (*r*), as well as *t*- and *p*-values, the latter of which was evaluated against an alpha level of 0.05 to determine correlation significance.

### Drift diffusion model

We fit hierarchical drift diffusion models to participants’ choice and reaction time data using the HDDM toolbox (version 0.9.8; Wiecki et al., 2013) in Python (version 3.7.12, van Rossum & Drake, 2009) via Jupyter Notebook (version 6.5.5; Kluyver et al., 2016). HDDM implements a hierarchical Bayesian estimation method, which allows for the simultaneous estimation of individual- and group-level parameters at different hierarchical stages by assuming that model parameters for individual participants are randomly drawn samples from the group-level distribution. This hierarchical approach has been shown to lead to more reliable parameter estimation, especially when less data is available or single-trial neural data is used for parameter estimation (Matzke et al., 2013; Wiecki et al., 2013). Additionally, the Bayesian method generates joint posterior distributions for all parameters, which not only provides an estimate for the most likely value of the given parameter, but also quantifies uncertainty in parameter estimation.

Drift diffusion modelling (DDM; Ratcliff 1978) assumes that during two-choice decisions, responses are determined by a noisy evidence accumulation process. Evidence in favour of a choice alternative can fluctuate from time point to time point based on the level of stimulus noise, its neural representations, or the attention paid to task and stimulus features. Observed choice and RT data *x_p,i_* for participant *p* on trial *i* is represented by the DDM likelihood function (Navarro & Fuss, 2009);

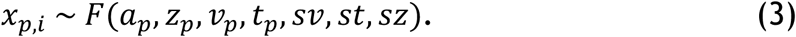

The rate of evidence accumulation is determined by the drift rate parameter *v* and specifies the amount of evidence accumulated per unit of time. The drift rate can reflect participants’ task efficiency as well as task difficulty, such that higher values of *v* produce faster and more accurate responses. A response is executed when enough relative evidence is accumulated in favour of a choice option, i.e., when the amount of relative evidence in favour of a stimulus crosses the decision threshold *a*. This parameter controls the speed-accuracy trade-off (Ratcliff & Rouder, 1998; Zhang & Rowe, 2014) and reflects response caution; higher values of this free parameter lead to slower but more accurate responses. The non-decision time parameter *t* captures RT components linked to the perceptual encoding of stimuli and motor processing after a choice is selected. Consequently, this parameter is independent of evidence accumulation, and affects RTs without influencing accuracy. The starting point bias parameter *z* captures bias towards one of the choice alternatives and determines the start of the drift diffusion process relative to the two boundaries. Thus, values of *z* > 0.5 suggest an *a priori* bias towards the upper boundary and values of *z* < 0.5 imply a bias towards the lower boundary. In case of no bias (i.e., when there is an equal amount of evidence in favour of both choice options), evidence accumulation begins halfway between the two boundaries, with *z* = 0.5. In all our models, we implemented accuracy coding; the upper boundary reflected choices where the symbol with the higher reward probability was selected, whilst the lower boundary reflected responses where the symbol with the lower reward probability was chosen by participants. As customary with accuracy coding, we fixed *z* at 0.5 and thus assumed no starting point bias.

We implemented the ‘full’ version of the DDM (Ratcliff & McKoon, 2008; Ratcliff & Rouder, 1998), which includes inter-trial variability parameters *st* and *sv* to capture the variability in non-decision time and drift rate, and starting point, respectively. The full DDM has been shown to outperform the simpler version of the model as it is able to account for the behavioural phenomena that errors are either faster or slower than correct responses (Ratcliff & Rouder, 1998). We estimated only the group-level values for these three parameters as the influence of the individual inter-trial variability parameters is often modest or even absent, with a large dataset required for reliable estimation (Wiecki et al., 2013).

To estimate the influence of the participant-specific EEG feedback components on subsequent decision making, we utilised the regression functionality within the HDDM toolbox. This functionality allowed us to model the linear link between our neural predictors and DDM parameters on a trial-by-trial basis by incorporating single-trial EEG component discriminant amplitudes (*y_i,_* Eq.1) into the parameter estimation process. Similar to our EEG-informed pupil analysis, the early and late EEG components were broken down by feedback valence, yielding four components in total; early positive, early negative, late positive, late negative (Fig.2a). We estimated posterior distributions for DDM parameters as well as for regression coefficients specifying the degree to which model parameters are influenced by the trial-to-trial variations in the electrophysiological measures.

To test our hypothesis that increased feedback processing by the early and late systems following negative feedback reduced evidence accumulation in subsequent trials, we utilised the above four single-trial component amplitudes as predictors to explain parameter variability. Specifically, component discriminating amplitudes from trial *i*-1 were used to predict the drift rate and/or boundary separation on trial *i* according to

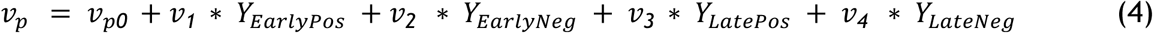

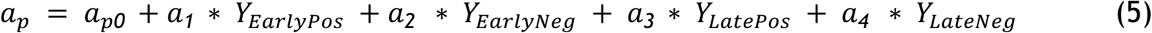

Here, *v_p0_* and *a_p_*_0_ represent the participant-specific intercept terms for the drift rate and boundary separation parameters, respectively. Additionally, v_1-4_ and a_1-4_ weigh the slope of the drift rate and boundary separation parameters, respectively, by the values of the four EEG component discriminant amplitudes on the previous trial. We utilised three separate neural models; either the drift rate (Model 1), boundary separation (Model 2), or both (Model 3) varied according to the four EEG component amplitudes. To more reliably compare the three different model variants, we estimated neural regression coefficients on the group-level only (Fig.1) as individual-level estimation of DDM regression coefficients has been suggested to increase parameter collinearity (Frank et al., 2015), and could therefore bias results.

To further probe the hypothesised relationship between LC-NA system mediated network resets in reward learning structures and reduced evidence accumulation in subsequent decisions, we adjusted the best-fitting neurally-informed DDM to estimate both individual- and group-level parameters for the neural regression coefficients. This in turn allowed us to explore the relationship between how the early or late negative components predict the pupil response during feedback and affect evidence accumulation on the following trial. We hypothesised that participants with a stronger coupling between the early/late negative component and the pupil response would also exhibit a stronger reduction in evidence accumulation in the next trial as a function of early/late system activity. To evaluate this hypothesis, we utilised robust bend correlations to quantify the relationship between the regression coefficients linking early/late negative discriminant amplitudes to the feedback-related pupil response and the regression coefficients determining the influence of the early/late negative component amplitudes on evidence accumulation in the next trial.

To evaluate our hypotheses regarding the impact of feedback processing on subsequent evidence accumulation and boundary separation, we utilised Bayesian hypothesis testing on the group-level DDM regression coefficients of the winning model. Carrying out significance tests directly on the model posteriors is a major advantage of the Bayesian model estimation method (Kruschke, 2010). We computed the proportion of the posterior distribution of the regression coefficients that were above or below 0. In case at least 95% of the proportion of the group-level coefficient posteriors were above (below) 0, we concluded that they had a significant positive (negative) association with the parameter of interest (Frank et al., 2015; Mattes et al., 2022; Weicki et al 2013).

For all our models, we utilised Markov Chain Monte Carlo (MCMC) sampling (Gamerman & Lopes, 2006) within the HDDM package to derive the joint posterior distribution for all parameters. For each model, we generated 10,000 samples, discarded the first 5,000 as burn-in, and thinned the remaining 5,000 samples by keeping every fifth draw, resulting in a total of 1,000 samples of the joint posterior distribution of the parameters. Following the recommendation by Wiecki and colleagues (2013), we used the default, informative priors implemented in the HDDM. These priors are based on results from a set of 23 studies, and were found to improve parameter recovery and reduce issues related to collinearity (Matzke & Wagenmakers, 2009). We used the default outlier setting in the HDDM, whereby 5% of RT outliers are removed to account for responses produced by processes other than drift diffusion, such as lapses in attention. Thus, we estimated a mixture model, whereby 5% of the trials are considered to be distributed according to a uniform distribution and the remaining 95% of trials are distributed according to the DDM likelihood function. This method of categorising each trial as a guess or a delayed startup has been found to improve parameter recovery in the presence of outliers (Frank et al., 2015; Verdonckhove & Tuerlinckx, 2007; Wiecki et al., 2013).

### Model convergence, selection, and predictive accuracy

To formally test model convergence, we ran four chains of each model and utilised the Gelman-Rubin 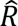 statistic (Gelman & Rubin, 1992) to compare within- and between-chain variances. Model convergence is commonly considered satisfactory when 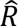 is close to 1 (when samples of the different chains are indistinguishable) but below 1.1 (Franzen et al., 2020; Pedersen & Frank, 2020; Pedersen et al., 2017). Additionally, we visually inspected all group-level sample traces, autocorrelation levels, and the marginal posteriors to further ensure appropriate model convergence. As such, we watched out for signs of poor convergence, including the presence of drifts and major jumps in the posterior sample traces, high autocorrelation levels that surpass the recommended level of 50 for any of the parameters (Wiecki et al., 2013), and non-normal marginal posterior distributions.

We used the Deviance Information Criterion (DIC; Spiegelhalter et al., 2002) to compare our models, as other measures, such as the Akaike Information Criterion (Akaike, 1978) or Bayesian Information Criterion (Schwarz, 1978) are not appropriate for hierarchical model comparison. The DIC is widely used for assessing the fit of hierarchical models by selecting the model that achieves the most optimal trade-off between model complexity and goodness-of-fit (Spiegelhalter et al., 2002). Lower DIC values are preferred as they are linked to models with the highest likelihood and the least degrees of freedom.

To evaluate the ability of our models to reproduce key elements of the data, we carried out posterior predictive checks. Specifically, we simulated 500 data sets from the posterior distribution of the parameters of the best fitting neurally-informed model, and compared the distance between the summary statistics of the predicted and observed data. By simulating multiple data sets from the posterior, we derived both summary statistics and a distribution of the simulated data, allowing us to capture uncertainty in our model estimates. The application of the DDM to reward learning tasks entails that the model should capture trends in both choice probabilities and RT distributions.

To examine RT trends across simulated and observed data, we plotted observed and predicted RT probability distributions alongside each other (Fig.5a). To further probe that simulated RTs and choices captured key trends in our data, we generated quintile probability plots (Ratcliff & McKoon, 2008; Frank et al., 2015). Fig.5b shows the mean observed and simulated RTs for each quintile (10th, 30th, 50th, 70th, and 90th) separately for the upper and lower boundaries. The shaded areas reveal estimation uncertainty as it represents predicted RTs within one standard deviation from the mean.

### Code accessibility

Linear discriminant analysis scripts are openly available at https://github.com/liinc-lab/EEGLAB-plugin-Linear-Discrimination.

## Results

### Probabilistic reversal-learning task performance

During the 300 trials of the experiment, participants reached a mean accuracy level of 66.89% (*SD* = 7.07%), encountered an average of 9.96, (*SD* = 0.20) reversals, and experienced *M* = 168.3 (*SD* = 10.58) positive and *M* = 128.27 (*SD* = 10.52) negative feedback trials. In line with reinforcement learning, participant’s choice behaviour appropriately tracked reversals (Fig. 2B); mean accuracy on the trial before reversals reached 68.93% (*SD* = 16.48%), dropped to 32.82% (*SD* = 15.83) on the trial where reversals were introduced, and plateaued on the 9th trial following reversals at 73.11 % (*SD* = 13.34%). Feedback type also significantly influenced choice behaviour; accuracy was higher (*t*(47) = 38.13, *p* < 0.001) following positive (*M*: 81.84%, *SD* = 5.51%) than negative (*M*: 47.51%, *SD* = 8.59%) feedback (Fig.2C), whilst participants were more likely to switch to the other symbol (*t*(47) = 12.81, *p* < 0.001) following negative (*M*: 44.07%, *SD* = 20.82%) than positive (*M*: 44.07%, *SD* = 20.82%) feedback trials (Fig.2E). There was no significant difference (*t*(47) = −1.02, *p* = 0.31) between the RTs in trials leading to positive (*M* = 614.42 ms, *SD* = 81.55 ms) and negative (*M* = 617.41 ms, *SD* = 87.13 ms) feedback (Fig.2D). Given that the above analyses confirmed that participants’ behaved in accordance with basic principles of reinforcement learning, we fit a Q-learning model with a dynamic learning rate to choice data. The model provided an excellent fit to our data (Fig.2F) and as anticipated, the direction of value-updating matched the valence of previous feedback, i.e., the value of the chosen symbol increased (decreased) following positive (negative) feedback (Fig.2G).

### Multivariate EEG analysis

To establish temporally distinct neuronal components linked to feedback valence, we used single-trial multivariate discriminant analysis of feedback-locked EEG signal (Parra et al., 2005). Specifically, using a sliding window approach, we estimated linear weights of the multi-channel EEG signals that achieved maximal discrimination between positive and negative trials within each time window. We repeated this analysis separately for each participant. Projecting the original data through these linear weights produced an aggregate representation of the discriminating component amplitude (*y_i_*, Eq.1) for each trial, which in turn we used as a proxy of how the participants encoded the feedback on individual trials. These discriminant component amplitudes reflect the distance of each trial from the discriminating hyperplane and here can be interpreted as a proxy for the neural variability elicited by positive and negative feedback, with the shared signal variance removed (Fouragnan et al., 2015; 2017; Franzen et al., 2020; Parra et al., 2005, Sajda et al., 2009). To assess the performance of the discriminator, we estimated the area under a receiver operating characteristic (ROC) curve combined with a leave-one-trial-out cross-validation method for each participant (Philiastides & Sajda, 2006; Philiastides, Ratcliff, & Sajda, 2006). Due to the linearity of our model, we also derived scalp topographies (Eq.2) representing the electrical coupling between the discriminating component amplitudes and the observed EEG signal.

Population discriminator performance over time and the scalp topographies associated with the most prominent peak timepoints are shown in Figure 5a. Consistent with previous work (Fouragnan et al., 2015; 2017), the discriminator reached significance between 150 and 590 ms after feedback presentation. We found a clear transition of topographies around 300-320 ms post-feedback, suggestive of two separate systems unfolding sequentially in time as in previous work (Fig. 5a). To identify participant-specific early and late feedback valence components, we selected peak discrimination times within two pre-defined time windows separated by the transition point in the group topographies as outlined above (i.e. 150-290 ms and 300-550 ms post-feedback for the early and late components, respectively).

Using this approach, we identified the same broad and distinct spatial profiles as previously established for the early and the late valence components. The early component peaked, on average, at 227 ms (*SD* = 36 ms) following feedback presentation with an *A_z_* value of .68 (*SD* = .07) and a corresponding scalp topography reflecting a midline cluster, mainly over central and centroparietal electrode sites. The late component peaked, on average, at 414 ms (*SD* = 75 ms) following feedback presentation with an average *A_z_* value of .70 (*SD* = .07) and a corresponding scalp topography showing lateral temporoparietal electrode clusters as well as a broad midline cluster opposite in polarity compared to the early component. Both components share a strong temporal and spatial resemblance to the analogous components observed in previous studies (Carvalheiro & Philiastides, 2023; Fouragnan et al., 2015; 2017; 2018; Westwood & Philiastides, 2024).

To verify that the early and late components reflect a valence effect, rather than signed RPEs, we divided the single-trial discriminator component amplitudes (*y_i_*) into four bins according to the value of the RPEs estimated by our reinforcement learning model (i.e. high negative, low negative, low positive, high positive). Similar to the results by Fouragnan et al. (2015; 2017), both components revealed a profile consistent with a categorical valence effect (with overall higher discriminator values for positive compared to negative feedback), with no additional modulation by RPE magnitude. We also formally compared the single-trial discriminator component amplitudes with the trial-wise RPE estimates from the model and found low within-participant robust bend correlation coefficients for both the early (positive RPE: *r* = −0.03, negative RPE: *r* = - 0.002) and the late (positive RPE: *r* = −0.04, negative RPE: *r* = −0.01) components.

In all subsequent analyses, we leverage the participant-specific discriminator component amplitudes (*y_i_*, Eq.1) linked to the early and late components as an index of how the valence of each feedback is encoded in individual trials across time (Fouragnan et al., 2015; 2017; Franzen et al., 2020; Westwood & Philiastodes, 2024). Importantly, trial-wise amplitude variations between the early and late components were generally uncorrelated as revealed by robust bend correlation coefficients (*r* = 0.08 for positive feedback, *r* = 0.12 for negative feedback). This allowed us to build parametric EEG-informed regressors independently for each of the two temporal components to predict behavioural and pupil data.

### Pupil data analysis

The population pupil response peaked at around 1000 ms post-feedback, after which it plummeted at around 2200 ms, followed by a modest rise. This is compatible with previous findings indicating peak pupil dilation around 1000 ms following stimulus onset (van Rij et al., 2019). Prior to exploring the trial-wise covariation in pupil and EEG dynamics, we examined the overall pupil response to positive and negative feedback (Fig.4A). For this, we utilised non-parametric cluster-based permutation *t*-tests (Maris & Oostenveld, 2007) for dependent samples to compared the mean response per feedback type for each 25 ms time window from the 500 ms before the onset of feedback presentation until 3000 ms post-feedback. Pupil dilation was significantly different between the feedback types in the 225-3000 ms post-feedback interval (*p* < 0.001).

**Figure 4.**
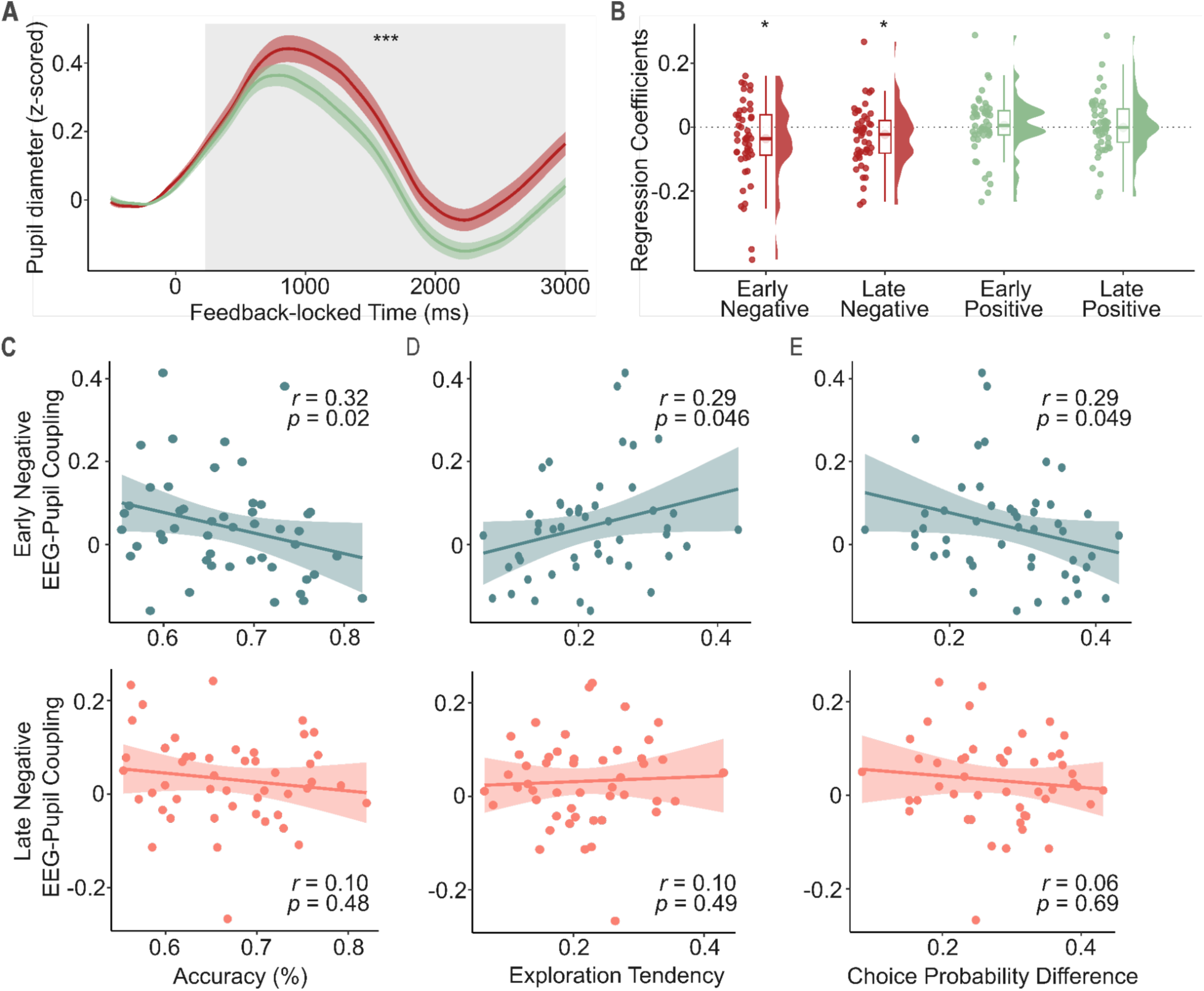
Pupil data analyses. **A.** Population pupil response linked to positive and negative feedback trials. The grey shaded area corresponds to the period of time (225-3000 ms) with a significantly different (*p* < 0.001) feedback-evoked pupil response for negative versus positive feedback trials, evaluated by a non-parametric cluster-based permutation *t*-test. **B.** Results of our GLM predicting the full course pupil data using the four EEG-derived predictors (early positive, early negative, late positive, late negative). *T*-tests revealed that unlike the early and late positive, the early and late negative component amplitudes significantly predicted (*p* < 0.05) the pupil response. **C.** Across-participant correlation between the early (top) and late (bottom) negative EEG-pupil coupling and mean accuracy. **D.** Across-participant correlation between the early (top) and late (bottom) negative EEG-pupil coupling and absolute choice probability difference. **E.** Across-participant correlation between the early (top) and late (bottom) negative EEG-pupil coupling and exploration tendency (i.e., the proportion of choices in which the symbol with the lower reward probability was selected). For **C-E**, the sign related to the EEG-pupil coupling (i.e., the regression coefficients from the EEG-pupil GLM) was flipped to support easier comprehension of the relationship between coupling strength and behavioural metrics.

This feedback valence-induced difference in pupil dilation is consistent with our hypothesis that negative feedback prompts elevated LC-NA system activity reflecting increased levels of contextual uncertainty and a neural interrupt signal. However, this comparison of population-level pupil responses provides no insight into whether this valence-related difference in pupil response is a general arousal effect induced by an increased response to negative feedback, a reduced reaction to positive feedback, or both. To resolve this issue and unmask latent trends in the data potentially concealed by averaging across trials, we utilised the single-trial variability in EEG component discriminant amplitudes to predict the pupil response.

### EEG-informed pupil analysis

To test the hypothesised link between the two feedback EEG-components and the pupil response, we exploited the trial-wise EEG component amplitude fluctuations within each component type (early and late), and independently for positive and negative feedback, to predict the full course pupil response within a general linear modelling framework. We thus built four separate trial-wise EEG predictors; early positive, early negative, late positive, and late negative which allowed us to establish EEG component-pupil coupling effects beyond any added unspecific effects of overall feedback valence. To further account for the possible effect of feedback surprise/salience as well as nonspecific effects elicited by the main task events, we constructed an extended model. This more complex model included a separate regressor based on the unsigned RPEs estimated by our reinforcement learning model as well as three unmodulated boxcar regressors at the time of choice stimuli onset, choice, and feedback onset (Fig.1A). All predictors in both models were convolved with the participant-specific PuRF.

The model with the additional predictors provided a better fit to the pupil data compared to the simpler model with the EEG-only predictors (RMSE of 0.98 and 1, respectively). Consequently, we used the regression coefficients linked to the neural regressors derived from the more complex model in all our subsequent analyses. In line with our hypotheses, the early (*t*(47) = −2.37, *p* = 0.02) and late negative (*t*(47) = −2.36, *p* = 0.02), but not the early (*t*(47) = .38, *p* = 0.70) and late positive (*t*(47) = .35, *p* = 0.72) EEG component amplitudes significantly predicted the pupil response (Fig.4B).

Importantly, more negative early and late negative component amplitudes correspond to boosted neural reactivity to negative feedback. Therefore, the negative association between the early and late negative component amplitudes and the pupil response suggests that stronger neural encoding of negative feedback results in a more pronounced pupil response. It is noteworthy to remember that we found no significant difference between the early and late negative component coefficients (*t*(47) = −0.49, *p* = 0.63), suggesting that the two components may represent similar neural processes. This is in line with our initial hypotheses that the early system could be guided by the LC-NA arousal network in response to negative feedback whilst exhibiting a downstream influence on the late system in order to down-regulate value-encoding.

Our mixed effects linear model (see Methods) further confirmed these results; the early (*t*(47) = −3.39, *p* < 0.001) and late negative (*t*(47) = −2.44, *p* = 0.01) component amplitudes significantly predicted the feedback-evoked pupil response, unlike the early (*t*(47) = 1.28, *p* = 0.20) and late positive (*t*(47) = 0.24, *p* = 0.81) component amplitudes.

In line with the proposed role of the LC-NA system in signalling unexpected uncertainty and implementing network resets (Bouret & Sara, 2005; Yu & Dayan, 2005), we expected that diminished performance and increased uncertainty regarding which option to choose would increase the likelihood of initiating such interrupt signals. To formally test whether a stronger coupling between the early system and the pupil response following negative feedback reflects such network resets, we correlated the participant-specific EEG-pupil coupling (i.e.,the early and late negative component regressions coefficients derived from the EEG-informed pupil GLM) with accuracy, uncertainty, and exploration tendency.

We found a significant correlation between task accuracy and the early (*r* = 0.32, *p* = 0.02), but not the late (*r* = 0.10, *p* = 0.48) EEG-pupil coupling (Fig.4C). In addition, the early (*r* = 0.29, *p* = 0.046) but not the late (*r* = 0.10, *p* = 0.49) EEG-pupil coupling significantly correlated with difference in overall choice probabilities between the two symbols (Fig.4C). The strong correlation between participant-specific mean accuracy and choice probability difference (*r* = 0.80, *p* < 0.001) further indicates that participants who experienced less uncertainty regarding which symbol to choose performed better. Overall, these results suggest that lower accuracy levels and increased choice uncertainty are linked to a stronger coupling between the feedback-related pupil response and negative feedback processing by the early, but not the late, system.

Uncertainty-induced network resets may also facilitate exploratory behaviour in order to support the establishment of a new model of the external world. Accordingly, we found that the increased EEG-pupil coupling for the early (*r* = −0.29, *p* = 0.049), but not the late (*r* = −0.06, *p* = 0.67), negative component was linked to an increased propensity to choose the symbol with the lower reward value (Fig.4C). Correlation coefficients of similar magnitude (*r* = 0.26, *p* = 0.08 and *r* = 0.03, *p* = 0.86 for the early and late negative components, respectively) were obtained when using the participant-wise inverse temperature parameter from our reinforcement learning model as a measure of exploration tendency. Lower values in the inverse temperature parameter index reduced sensitivity to reward deviations in the available options, and consequently, they mark an increased exploration propensity. We found a strong correlation (*r* = 0.90, *p* < 0.001) between the two measures of exploration tendency, suggesting they represent the same underlying construct. This increased exploration propensity linked to a stronger pupil and early system coupling further implies that the LC-related early system could contribute to increased exploratory behaviour.

### Drift diffusion modelling results

To examine the knock-on effects of our EEG-derived feedback components on subsequent choice dynamics, we implemented neurally-informed drift diffusion modelling. In line with the hypothesised role of the early system in facilitating LC-induced network resets and down-regulating late system activity, we expected increased early and late systems activation following negative feedback to reduce evidence accumulation on the next trial. In addition, we expected that an early-system-related arousal signal after negative feedback may also result in more response caution and hence increased boundary separation.

To test these hypotheses, we estimated three different neurally-informed regression models within the DDM framework; we either varied the drift rate, boundary separation, or both according to our four single-trial EEG component (early positive, early negative, late positive, late negative) discriminant amplitudes (*y_i_*, Eq.1). The model with both drift rate and threshold varying dynamically as a function of our EEG component amplitudes (Model 3) achieved a better fit (DIC = 1938) compared to the model with only the drift rate (Model 1; DIC = 1953) or boundary separation (Model 2; DIC = 2210) varying as a function of the component amplitudes. To ensure that our models converged, we ran four chains of each model and utilised the Gelman-Rubin 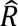 statistic (Gelman & Rubin, 1992) to compare within- and between-chain variances. All three models appeared to converge appropriately, with all parameter 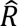 values lying under the recommended value of 1.1 (Franzen et al., 2020; Pedersen & Frank, 2020; Pedersen et al., 2017). Additionally, all parameter 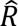 values of the best-fitting model were below 1.01, suggesting excellent model convergence. Group-level parameter estimates for the best fitting model are shown in Table S1.

To evaluate that the best-fitting model correctly captured RT and choice trends in our data, we simulated 500 sets of RT and choice data by drawing samples from the joint posterior distribution of model parameters for each participant. Predicted RT data for the higher- and lower-valued options were then pooled together from individual participants and plotted alongside the experimental data as posterior model predictions (Fig.5a; Frank et al., 2015; Franzen et al., 2020). Predicted and observed RTs followed a similar trend for both response types, implying that the model provided a good fit to the experimental data. To further examine model predictions for RT and choice data, we generated quintile-probability plots (Frank et al., 2015; Ratcliff & McKoon, 2008) depicting RT quintiles separately for each response type (Fig.5b). In all cases, observed RT quintiles were within one standard deviation of the predicted RT quintiles, confirming that our best fitting model provided a good fit to observed choice proportions and RT distributions.

**Figure 5.**
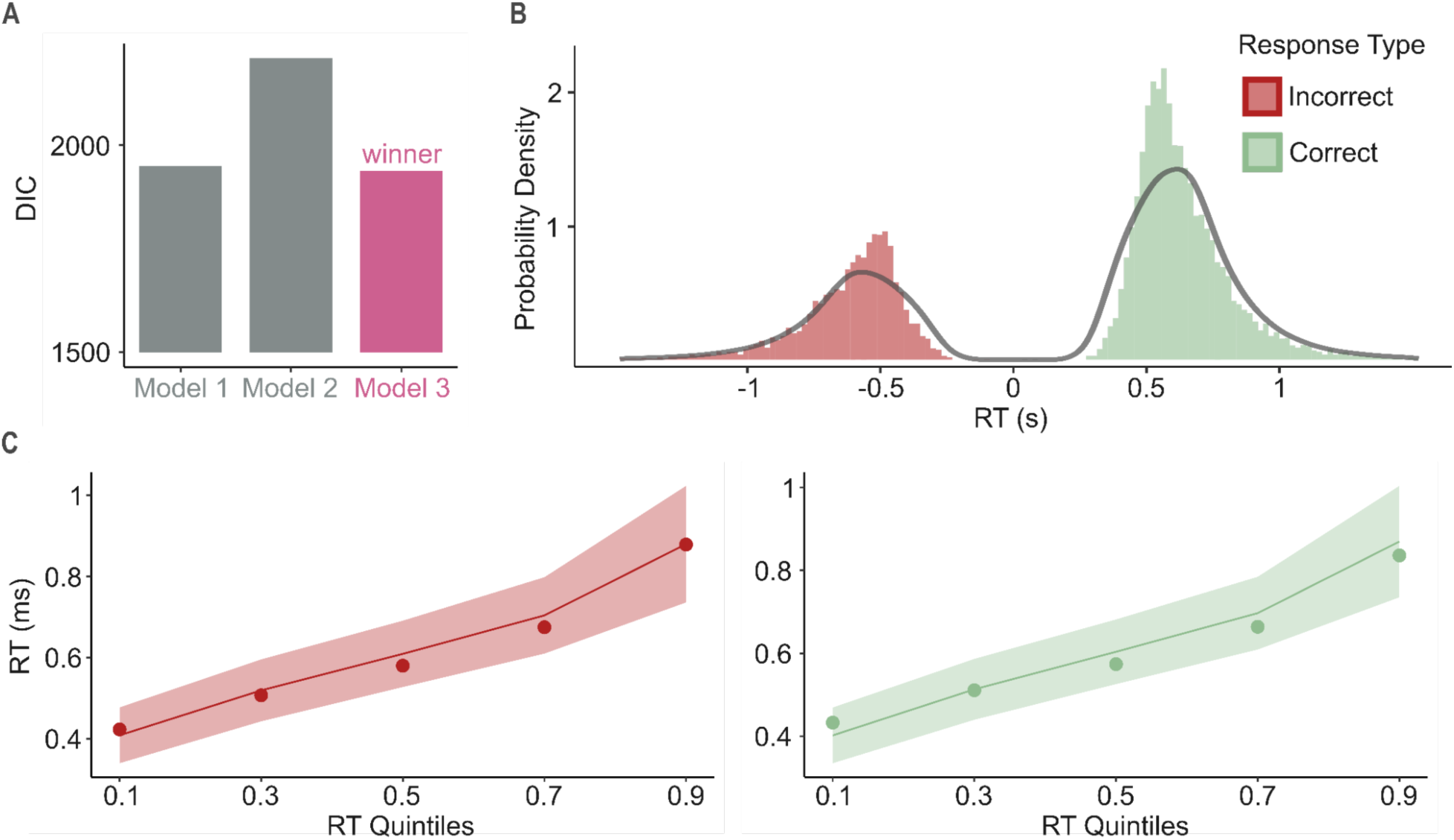
Model selection, observed and predicted RT distributions. **A.** We implemented three neurally-informed DDMs, where either the drift rate (Model 1), boundary separation (Model 2), or both (Model 3) varied according to our four trial-wise EEG feedback component discriminant amplitudes. Model 3 achieved a better model fit (DIC = 1938) than Model 1 (DIC = 1953) or Model 2 (DIC = 2210). To evaluate whether Model 3 appropriately captured RT and choice trends in our data, we carried out posterior predictive checks by drawing 500 sets of RT and choice data from the joint posterior distribution. **B.** Probability density distributions are shown for the observed RTs (histogram) and the RTs predicted by the posterior predictive simulations (probability density curve) from the best-fitting DDM. RTs on the negative scale portray RTs for the lower-valued (i.e., ‘incorrect’) option (RTs were sign-flipped), whilst the RTs on the positive scale portray RTs for the higher-valued (‘correct’) option. The relative area under the histogram and probability density curve represent choice proportions, demonstrating a higher proportion of choices for the higher-compared to lower-valued option. **C.** Quintile probability plots towards the upper boundary reflecting the higher-valued option (left) and towards the lower boundary reflecting the lower-valued option (right). RT quintiles (10th, 30th, 50th, 70th, 90th) are shown on the *x*-axes, with their corresponding observed (dots) and model-predicted (line) values on the *y*-axes. Predicted data were simulated from the winning model.The shaded areas represent one standard deviation above and below the posterior predictive RT distributions and capture estimation uncertainty.

### EEG feedback components shape subsequent decision making

We next investigated whether trial-to-trial variability in feedback-locked EEG components predicts how decisional parameters are adjusted in future choices. We utilised Bayesian hypothesis testing (Frank et al., 2015; Mattes et al., 2022; Wiecki et al 2013) on the regression coefficients linked to the four EEG components of the best-fitting neurally-informed model (Model 3) to explore whether they significantly explained variation in the drift rate and boundary separation over the next trial. If at least 95% of the posterior distribution of a neural regression coefficient were above (below) 0, we concluded that there was a significant positive (negative) association between the component and the parameter of interest (Frank et al., 2015; Mattes et al., 2022; Wiecki et al 2013).

Increasingly positive (negative) component amplitudes indicate that feedback was more positively (negatively) encoded on a given trial. The entire (100%) posterior distribution of the regression coefficients estimating the impact of the EEG early and late negative components on evidence accumulation was positive. Thus, the more positively feedback was encoded by the two systems, the more evidence accumulation increased in the next trial. Conversely, increased negative feedback encoding by the early and late systems reduced evidence accumulation on the next trial (evidenced by 100% and 99% of the posterior distribution shifted away from 0 for the early and late negative components, respectively). Furthermore, examining the mean posterior estimates for the group-level regression coefficients suggested that positive feedback processing by the early (*M* = 0.26, *SD* = 0.03) and late (*M* = 0.29, *SD* = 0.03) systems had a larger impact on evidence accumulation compared to the effect of the early negative (*M* = 0.13, *SD* = 0.04) and late negative (*M* = 0.09, *SD* = 0.04) components (Fig.6A).

**Figure 6.**
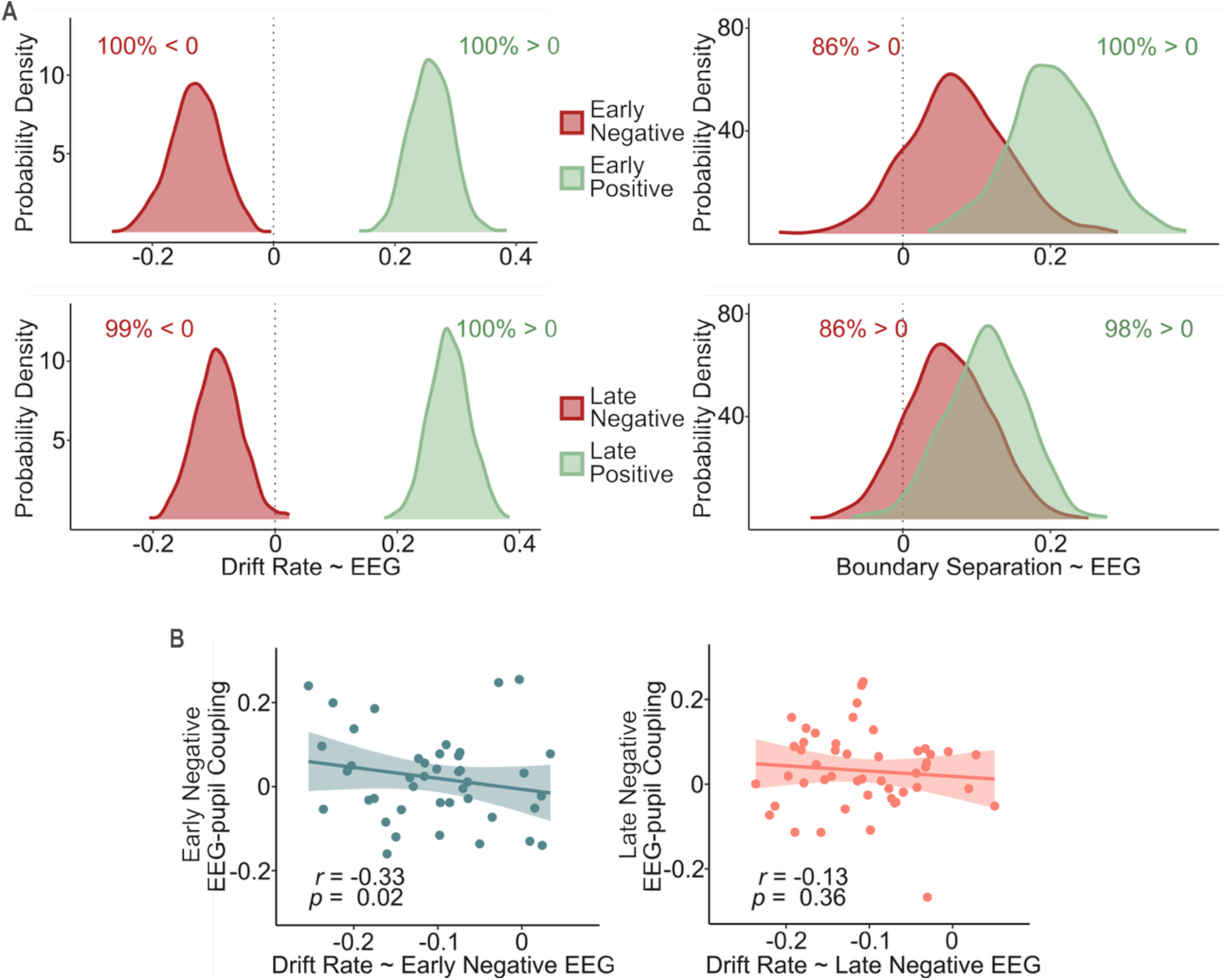
DDM results. **A.** Posterior distributions show the estimated effect of the four EEG component discriminant amplitudes on the drift rate (left) and boundary separation (right). Drift rate was significantly related to all four EEG components demonstrated by the posterior distributions being shifted away from zero. Boundary separation was significantly modulated by the early and late positive component amplitudes, but not by the early and late negative component amplitudes. The EEG component amplitudes explained a larger proportion of the variation in the drift rate compared to boundary separation, suggesting that the feedback-related accuracy difference following positive and negative feedback trials is primarily driven by drift rate fluctuations. Peak values of each distribution portray the best estimate for the relevant regression coefficient. The width of each distribution represents estimation uncertainty. The signs related to the early and late negative DDM regression coefficients were flipped in order to better visualise effect directions (i.e., increased negative feedback encoding was linked to reduced drift rates). **B.** Left: Across-participant bend correlation (r = −0.33, p = 0.02) between participants-specific early negative EEG-pupil coupling strength and individual-level DDM parameters linking the early negative component to evidence accumulation on the next trial. Right: Across-participant bend correlation (r = −0.13, p = 0.36) between participants-specific late negative EEG-pupil coupling strength and individual-level DDM parameters linking the late negative component to evidence accumulation on the next trial. The line and shaded area on each graph show the least-squares fit line and its 95% confidence band, respectively. The signs of the EEG-pupil and DDM regression coefficients were flipped in order to reflect that a stronger (more positive) early negative EEG-pupil coupling was associated with more significant (more negative) early negative component induced reductions in evidence accumulation.

There was strong evidence that decision threshold was associated with modulations in the early positive (*M* = .02, *SD* = 0.006, 100% of the posterior > 0) and late positive (*M* = 0.01, *SD* = 0.05, 98% of the posterior > 0) component amplitudes. In contrast, we did not find convincing evidence that the early (*M* = −0.007, *SD* = 0.07) or late (*M* = −0.006, *SD* = 0.06) negative components explained a significant proportion of the variation in this parameter (86% of the posterior < 0 for both the early and late negative components). Despite their lack of a significant effect on decision threshold, the early and late negative component regression coefficients suggest a similar influence; the more negatively feedback was encoded by either system, the more boundary separation increased on the next trial.

The EEG component amplitudes explained a larger proportion of the variation in the drift compared to boundary separation, suggesting that the difference in accuracy following positive versus negative feedback trials is mainly driven by fluctuations in the drift rate. Overall, boosted drift rates and boundary separation following increasing positive feedback encoding together with the reduced drift rates following negative trials explain the phenomenon that accuracy is higher following positive compared to negative feedback trials, whilst RTs remain similar.

To explore inter-individual differences in decision dynamics, we extended the above neurally-informed model to estimate not only group-but also individual-level parameters for the influence of our four EEG component discriminant amplitudes on subsequent evidence accumulation and boundary separation. Similar to previous models, convergence was evaluated by calculating 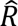 values based on four chains. All parameter 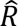 values (maximum: 1.04) were below the recommended value of 1.1 (Franzen et al., 2020; Pedersen & Frank, 2020; Pedersen et al., 2017), suggesting appropriate model convergence. The extended model achieved a better fit (*DIC*: 1856) compared to the model with the group-level-only estimates (*DIC*: 1938), and showed good predictive accuracy, with all quantiles of predicted data lying within the credible interval of the respective quantiles of the observed data. Reassuringly, Bayesian hypothesis tests on the group-level neural regression coefficients showed similar results to those derived from the previous model with group-level only estimates, suggesting reliable effects.

We predicted that across participants, a stronger early and late EEG-pupil coupling would be associated with larger reductions in evidence accumulation in response to negative feedback as both processes are hypothesised to reflect LC-NA mediated network resets in reward learning structures. To evaluate this proposition, we utilised across-participant robust bend correlations to quantify the link between the EEG-pupil coupling for the early (late) negative components and the DDM parameter estimates linking the early (late) negative component amplitudes to the evidence accumulation on the following trial. We found that the strength of the EEG-pupil coupling for the early (r = −0.33, p = 0.02), but not the late (r = −0.13, p = 0.36), component was significantly linked to drift rate reduction elicited by the trial-wise amplitude variation in the respective component (Fig.6B).

## Discussion

In this study, we exploited internal trial-by-trial variability in feedback-locked EEG signals during probabilistic reversal learning to validate the two spatiotemporally distinct neural systems associated with feedback evaluation (Fouragnan et al., 2015; 2017; Weiss et al., 2018). Crucially, we demonstrated that trial-wise amplitude fluctuations in both the early and late systems following negative feedback explained the differential feedback-valence effect observed in simultaneously acquired pupil responses. Moreover, increased early system EEG-pupil coupling was linked to elevated uncertainty estimation and exploration tendency, as well as reduced accuracy and evidence accumulation.

Whilst the relationship between pupil dilation and general arousal (salience) is well established (de Gee et al., 2014; 2017; 2021; Filipowicz et al., 2020; Preuschoff et al., 2011; Urai et al., 2017, Van Slooten et al., 2017; Varazzani et al., 2015), our results demonstrate that in addition to this broad unspecific effect, trial-wise variability in the early and late negative EEG component explains trial-wise changes in feedback-related pupil dynamics. This is suggestive of a systematic link between the LC-NA system and the distributed neural representations associated with these components (Algermissen et al 2024; Fouragnan et al., 2015), which could influence downstream choices and future behavioural adjustments.

Consistent with the finding that the early system down-regulates the late one following negative feedback (Fouragnan et al., 2015), we speculate that the differential coupling between the late system, likely under dopaminergic control (Fouragnan et al., 2015), and the pupil response following negative feedback reflects regulatory control originating in the early system. Accordingly, we observed that larger associations between early negative component amplitudes and the pupil response were negatively correlated with overall accuracy and positively correlated with choice ambiguity and exploration tendency. Furthermore, participants whose negative feedback encoding by the early system more strongly predicted value updating (reflecting increased internal estimates of uncertainty) were characterised by poorer performance.

These results are consistent with the LC-NA network reset hypothesis, which proposes that phasic burst of LC-NA activity promotes adaptive behaviour in response to unexpected events by transiently disrupting ongoing neural activity and reconfiguring task-relevant network dynamics (Bouret & Sara, 2005, Dayan & Yu, 2006; de Gee et al., 2017; Filipowicz et al., 2020; Sara, 2009; Urai et al., 2017). Our results support the proposition that increased estimates of contextual uncertainty, produced by negative feedback, are signalled to early system structures by increased LC-NA activity, in turn interrupting processing in the late network in an attempt to reduce the influence of learnt reward contingency representations. Such network resets would consequently decelerate subsequent decision making as evidence is accumulated in favour of a new, competing hypothesis (reversal), thereby increasing the likelihood of exploration (choosing the option that was previously considered inferior).

Accordingly, our neurally-informed drift diffusion model revealed that increased negative feedback processing reduced the drift rate on the next trial, without a significant change in boundary separation, in turn resulting in reduced accuracy. Increased positive feedback processing produced higher drift rates and boundary separation on the next trial, which contributed to improved accuracy. Across-participants, a stronger early EEG-pupil coupling was associated with larger early-component-induced reductions in evidence accumulation following negative feedback. These results further support the interpretation that the LC-NA-related early system decreases the influence of existing value representations in the late system, as reflected by the reduced evidence accumulation following increased negative feedback processing.

Our finding that increased positive feedback processing boosts boundary separation in subsequent trials contradicts results from previous studies. Within-trial decision conflict has been shown to increase boundary separation, presumably in an effort to buy more time for evidence accumulation during heightened decision difficulty (Cavanagh et al., 2011; Fontanesi et al., 2019; Frank et al., 2015). s positive feedback likely decreases decision conflict, it is expected to reduce boundary separation. However, unlike the above studies, we used a reversal learning paradigm and investigated inter-trial associations, which differences likely contribute to the conflicting results. Participants in our study could have increasingly expected reversals following positive feedback, in turn producing widening boundary separation. Future research based on reversal learning paradigms should confirm how positive feedback affects boundary separation in subsequent trials.

Consistent with our results and the network reset hypothesis, phasic LC-NA activity and the feedback-evoked pupil response have been linked to uncertainty signalling (Colizoli et al., 2018; Dayan & Yu, 2006; de Gee et al., 2014; 2017; Lavín et al., 2014; Pajkossy et al., 2017; Preuschoff et al., 2011; Urai et al., 2017; Yu & Dayan, 2005; van Slooten et al., 2017; 2018), exploration (Aston-Jones & Cohen; 2005; Gilzenrat et al., 2010; Jepma & Nieuwenhuis, 2011; Yu & Dayan, 2005; 2009; van Slooten et al., 2018), diminished accuracy and post-error slowing (Aston-Jones & Cohen, 2005; Murphy et al., 2016; Urai et al., 2017)., asl well as increased neural gain (Aston-Jones & Cohen, 2005; Bouret & Sara, 2005; Dayan & Yu, 2006; Eldar et al., 2013; Filipowicz et al., 2020; Gilzenrat et al., 2010; Krishnamurty et al., 2017; Nassar et al., 2012; Zenon, 2019). These results further indicate the LC-NA system in adaptively regulating the cortical influence of new versus learnt information.

For the LC to implement network resets, it needs to have access to a dynamic combination of bottom-up and top-down information (Filipowicz et al., 2020). The anterior cingulate cortex (ACC) is a prime candidate for providing these inputs to the LC (Aston-Jones & Cohen, 2005; de Gee et al., 2017) as it has strong reciprocal connection with the LC (Briand et al., 2007; Joshi & Gold, 2020), it has access to both top-down and bottom-up information via its robust connections with prefrontal structures and sensorimotor areas, and it is a prominent structure of the early system(Fouragnan et al., 2015).

ACC activity has also been found to mediate pupil responses (Joshi et al., 2016; Joshi & Gold, 2020; Murphy et al., 2014 and facilitate explore–exploit decisions (Fouragnan et al., 2019; Kolling et al., 2012). Chakroun et al., 2020; Daw et al., 2006; Forstmann et al., 2006; de A Marcelino et al., 2023). Furthermore, it promotes adaptive behaviour, including task difficulty monitoring (Ullsperger & von Cramon, 2001), conflict processing (Botvinick et al., 2001; 2004; Etkin et al., 2011), error detection (Rushworth et al., 2008; Yeung et al., 2004),and task volatility tracking (Behrens et al., 2007). Accordingly, adaptive gain theory (Aston-Jones & Cohen, 2005) proposes that ACC signalling to the LC produces phasic noradrenergic firing, which in turn increases cortical NA levels that enhance cognitive processing, perhaps by resetting reward learning networks. Further research is nevertheless needed to determine the precise role the ACC plays in facilitating network resets and influencing subsequent choices.

### Limitations and future directions

In this study, we examined reward learning in the appetitive domain. However, recent evidence suggests that despite the early and late components share similar behavioural and cortical signatures during approach and avoidance learning, LC activity is linked to the early component during negative feedback processing only in the appetitive domain (Carvalheiro & Philiastides, 2023), whilst increased pupil responses were reported following aversive compared to appetitive conditions (Finke et al., 2021; Westwood & Philiastides, 2024). Thus, the early and late components appear to be fundamental constituents of the reward learning process, further research is required to disassociate the unique contribution of the LC-NA network under different learning conditions.

Despite considerable evidence indicates the LC in generating the changes in pupil size related to cognitive processes (Joshi et al., 2016; Joshi & Gold, 2020; Murphy et al., 2014; Reimer et al., 2016), other brainstem nuclei, including dopaminergic, cholinergic, and serotonergic structures, as well as their connections with the LC, could partly moderate this pupil effect (de Gee et al., 2017; Reimer et al., 2016; Urai et al., 2017; van Slooten et al., 2018). Similarly, different neurotransmitters have been associated with adaptive learning processes; dopamine and acetylcholine have been linked to explore-exploit mechanisms (Chakroun et al., 2020; Cinotti et al., 2019; Frank et al., 2009; van Slooten et al., 2019)and and expected uncertainty signalling (Yu & Dayan, 2005), respectively.

Thus, both pupil dilation and the rapid network reconfigurations needed for adaptive learning are most likely achieved by different neurotransmitter systems operating in concert. Further research is needed to disentangle the complementary contribution and mode of action (i.e., phasic vs tonic) of each neurotransmitter system to these processes. Whilst pupillometry is an inexpensive and non-invasive tool for such investigation, high-resolution imaging of brainstem nuclei, high-precision causal manipulation (Murphy & Fouragnan, 2024), or single-cell recordings in animals could provide particularly influential insights.

### Conclusion

Utilising single-trial electrophysiological, pupillometry, behavioural, and modelling approaches, our study linked the two known spatiotemporally distinct reward learning components (Algermissen et al., 2024; Fouragnan et al., 2015) to the LC-NA system and its proposed role in uncertainty processing and network resets during reward learning. Our EEG- and pupillometry-based research methods could also provide valuable insights about altered neural mechanisms in neuropathological conditions characterised by altered noradrenergic activity, including dementia (Herrmann et al., 2004), mood and anxiety disorders (Brunello et al., 2003; Ehlers & Todd, 2017; Leonard, 1997), or addiction (Torregrossa, 2019).

## Acknowledgements

This work was supported by a European Research Council consolidator grant (DyNeRfusion; 865003 to M.G.P.) and the Economic and Social Research Council (ES/L012995/1 to M.G.P.). E.F. is funded by a UKRI FLF (MR/Y034368/1), a BBSRC (BB/Y001494/1), a Neuromod+ grant (EP/W035057/1) and an ARIA grant (SCNI-PR01-P15). K.B. was supported by the University of Glasgow MVLS doctoral training programme.

## Author Contributions

M.G.P. & E.F. designed the research. M.G.P. & K.B. performed the research and analysed the data. M.G.P, E.F., & K.B. wrote the paper.

## Data Availability

Data related to the manuscript are available upon request from the corresponding authors.

## Supplemental Results

### Analysis of EEG and behavioural data

To investigate the link between our EEG components and behavioural measures, we carried out four separate regression analyses using the single-trial early and late components discriminant amplitudes (*y_i_,* Eq.1) broken down by feedback type (positive and negative). Thus, each participant-specific regression analysis included four predictors; early positive, early negative, late positive, and late negative, the values of which were determined by the trial-wise linear discriminant amplitudes differentiating between positive and negative feedback. Higher discriminant amplitudes (more positive values) of the early and late positive components correspond to increased neural reactivity to positive feedback, whilst lower values indicate reduced encoding of positive feedback. Conversely, lower discriminant amplitudes (more negative values) of the early and late negative components correspond to boosted neural reactivity to negative feedback, whilst higher values indicate decreased encoding of negative feedback. By breaking down the feedback-related EEG response into an early and a late component for each feedback type, our four parametric EEG predictors utilised in the above regression analyses effectively removed overall valence effects (Westwood & Philiastides, 2024).

First, we performed a binomial logistic regression with a probit link function to determine whether the four single-trial EEG components on each trial are predictive of participants’ switching behaviour (i.e., a binomial variable indicating whether the participant chose the other symbol on the following trial). Our results (Supplementary Fig.1A) suggest that all four components predict switching behaviour (*t*(47) = −7.80, *p* < 0.001 for early positive, *t*(47) = −5.20, *p* < 0.001 for early negative, *t*(47) = −9.21, *p* < 0.001 for late positive, and *t*(47) = −3.92, *p* < 0.001 for late negative). In line with the findings of Fouragnan and colleagues (2015), the more positive decision feedback were neurally encoded, suggested by higher positive discrimination amplitudes, the probability of selecting the same symbol over the next trial increased. Similarly, the more negatively the decision feedback was neurally encoded, as reflected by more negative discrimination amplitudes, the more probability of selecting the same symbol over the next trial decreased. These findings imply a valence-specific component influence, whereby positive feedback processing depresses and negative feedback processing promotes avoidance learning.

Next, a binomial logistic regression tested whether any of the four EEG components predicted participants’ explorative choices, that is, whether they chose the symbol with the lower associated value as determined by our reinforcement learning model (Daw et al., 2006; Harada, 2020; Warren et al., 2017). Our results (Supplementary Fig.1B) showed that higher early positive (*t*(47) = −7.80, *p* < 0.001) and the late positive (*t*(47) = −5.20, *p* < 0.001) component amplitudes significantly reduced the likelihood of choosing the symbol with the lower value. At the same time, increased negative feedback encoding by the early (*t*(47) = −5.20, *p* = 0.03) and late (*t*(47) = −5.20, *p* = 0.01) systems increased participants’ inclination to explore over the next trial. This is consistent with the hypothesis that increased negative feedback encoding, reflected by more negative discriminant amplitudes, could promote an LC-induced cortical reset in reward learning structures, potentially linked to variability in the early system, which signals reversals (i.e., unexpected uncertainty arising from negative feedback), and in turn promote exploratory behaviours appropriate for establishing a new model of the prevailing reward contingencies.

Additionally, we investigated how our EEG component amplitudes affected the value updating process. To quantify trial-wise value-updating, we utilised the value difference of the chosen symbol between the current and next trials as derived from our reinforcement learning model. Supplementary Fig.1C depicts the participant-specific linear regression coefficients by predictor type. All four EEG components were highly predictive of value updating (*t*(47) = 12.50, *p* < 0.001 for early positive, *t*(47) = 4.31, *p* < 0.001 for early negative,*t*(47) = 12.67, *p* < 0.001 for late positive, and *t*(47) = 5.59, *p* < 0.001 for late negative). Similarly to Fouragnan et al. (2015), we found a positive relationship between the extent of up-and downregulation of value and the amplitudes of the EEG components. Accordingly, more positive component EEG discrimination amplitudes were associated with an increasing likelihood that the value of the chosen symbol would increase compared to the previous trial. Similarly, more negative EEG component discrimination amplitudes were linked to a higher likelihood that the value of the chosen symbol would decrease compared to the previous trial.

Stronger across-participant associations between value updating and the early (*r* = -.50, *p* < 0.001) or the late (*r* = -.61*, p* < 0.001) negative predictor amplitudes were linked to lower accuracy levels, as assessed by robust bend correlations (see Methods). The degree to which early (*r* = −0.10, *p* = 0.52) or late (*r* = −0.03, *p* = 0.84) positive component amplitudes predicted value updating had no significant correlation with accuracy. Specifically, the more participants’ negative feedback encoding predicted value updating, the worse they performed in the task. Due to the probabilistic nature of the task, participants occasionally received negative feedback despite having selected the symbol with the higher reward probability (expected uncertainty), rendering significant value adjustments unnecessary. Nonetheless, negative feedback could also signal a reversal of reward contingencies (unexpected uncertainty), in which case larger alterations in value representations are adaptive. Thus, the more strongly participants associate negative feedback with a reversal, the larger adjustments in value representations are expected. Consequently, our results are consistent with the interpretation that a more robust link between negative feedback processing and value updating reflects the overinterpretation of negative feedback (suboptimally large internal estimates of unexpected uncertainty), which in turn results in lower performance. These results are consistent with the hypothesis that an LC-NA related early system, which down-regulates the late system following negative feedback, signals unexpected uncertainty.

Our last, linear regression assessed whether our EEG component amplitudes predicted response caution, indexed by the difference in the *z*-scored reaction times (RTs) between consecutive trials. As Supplementary Fig.1D shows, only the early component following negative feedback was predictive of response time change (*t*(47) = −2.05, *p* = 0.046). The early positive (*t*(47) = 1.76, *p* = 0.085), late positive (*t*(47) = 0.73, *p* = 0.47), and late negative (*t*(47) = 0.66, *p* = 0.51) component amplitudes showed no significant association with response caution. This suggests that the more strongly negative feedback is encoded by the early system, the more response caution increases.

This result matches the proposed function of the early system in network resets, which in turn could slow down evidence accumulation over the next trial as evidence in favour of a new hypothesis (i.e., reversal) is accumulated.

**Supplementary Figure 1.**
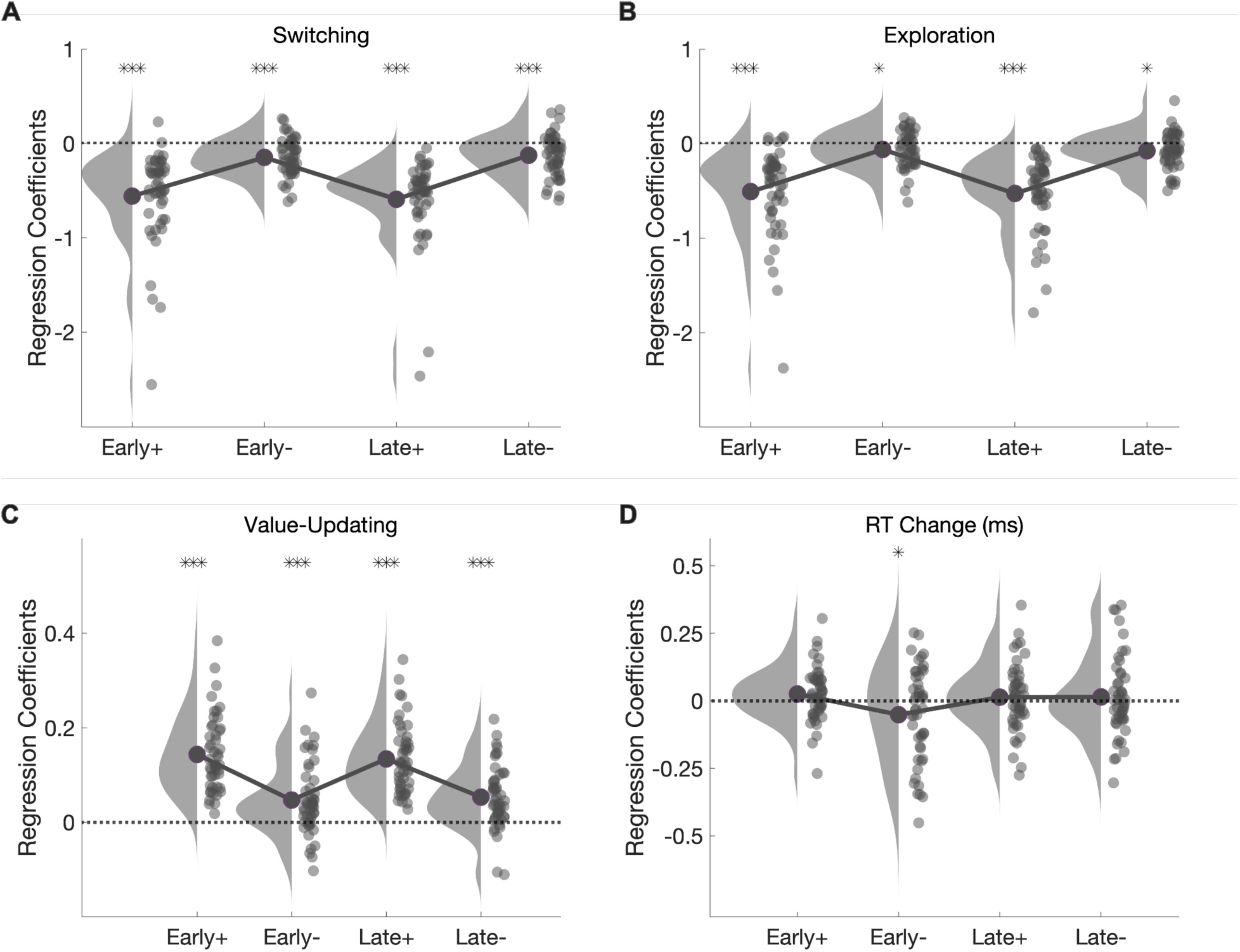
Single-trial EEG component amplitudes predict behaviour. **A,** Results of the binomial logistic regression, using four EEG-derived regressors at trial *t*, predicting the probability of switching at trial *t*+1. *T*-tests indicate all four components to be significant predictors of switching (*p* < 0.001 in all cases). **B,** Results of the binomial regression, using four EEG-derived regressors at trial *t*, predicting exploration (choosing the lower-valued option) over the next trial. Increasing early and late positive component amplitudes were associated with decreased explorative choices over the next trial (*p* < 0.001), whilst increased negative feedback encoding by the early (*p* = 0.03) and late (*p* = 0.01) systems boosted exploration on the next trial. **C,** Results of the linear regression, using the four EEG-derived regressors at trial *t*, predicting value updating at trial *t*+1. *T*-tests indicate all four components to be significant predictors of value updating (*p* < 0.001 in all cases). **D,** Results of the linear regression, using four EEG-derived regressors at trial *t*, predicting response caution between consecutive trials. *T*-tests indicate that only the early component linked to negative feedback is a significant predictor of response caution (*p* = 0.046), unlike the early component associated with positive feedback (*p* = 0.09), or the late component linked to positive (*p* = 0.47) or negative (*p* = 0.51) feedback.

### Behavioural-only drift diffusion model

To verify our results from the best fitting neurally-informed model (Model 3), we fit a DDM to our data with a behavioural-only feedback type regressor. Specifically, we modelled the drift rate and boundary separation as a linear function of a binary predictor specifying whether the previous feedback was negative according to

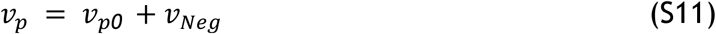

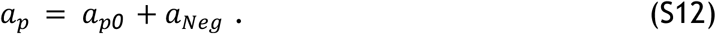

Here, *v_p0_* and *a_p0_* represent the participant-wise drift rate and boundary separation, respectively, following positive feedback trials, whilst *v_Neg_* and *a_Neg_* represent the effect of negative feedback on the drift rate and boundary separation, respectively. Similarly to Model 3, we included the inter-trial variability parameters *st* and *sv* in our model and assumed no starting point bias (*z* fixed at 0.5). Moreover, we estimated inter-trial variability parameters and the effect of negative feedback on the drift rate and boundary separation on the group-level only. All other parameters were derived on both the group- and individual-levels. We expected our results from the behavioural-only and neurally-informed models to align, with a similar effect of feedback type/encoding on the drift rate and boundary separation parameters.

The behavioural-only model successfully converged, with all parameter 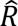 values below 1.1 (maximum: 1.004) and with a DIC value of 1679. Posterior predictive checks revealed that the model provided a satisfactory fit to our data, with model-predicted and observed RT and choice trends following a similar pattern. Consistent with the results from Model 3, Bayesian hypothesis testing on the group-level regression coefficients showed that negative feedback decreased the drift rate (*M* = −0.99, *SD* = 0.05, 100% of the posterior < 0) and increased boundary separation (*M* = −0.05, *SD* = 0.0008, 100% of the posterior < 0) on the next trial.

Crucially, results from our neurally-informed and behavioural-only models converged in supporting our hypothesis that (increased) negative feedback (processing by the early and late systems) reduce evidence accumulation on the next trial, suggesting the reliability of this effect. Further consistent with Model 3, (increased) positive feedback (processing by the early and late systems) increased both boundary separation and the drift rate in the next trial.

However, the two models produced contradictory results regarding the modulatory effect of negative feedback (processing) on boundary separation in the next trial. Whilst the neurally-informed model showed that the strength of negative feedback processing insignificantly increased decision threshold in the following trial, the behavioural-only model indicated negative feedback to reduce boundary separation in the successive trial. Previous results are consistent with the neurally-informed model as they found that the post-error response slowing generated by negative feedback resulted from an increased response threshold (Dutilh et al., 2012; Goldfarb et al., 2012).

The inability of the behavioural-only model to account for this effect (model falsification) is ground for absolute model rejection (Palminteri et al., 2017). We speculate that the conflicting results from the neurally-informed and behavioural-only models are introduced by the difference in the amount of information the neural and behavioural predictors carry. Neural constraints can account for a significant proportion of noise in cognitive models and ignoring these trial-to-trial variations in neural activity can lead to the overestimation of noise in the decision process (Franzen et al., 2020; Nunez et al., 2017; Turner et al., 2015). The inclusion of neural predictors in our model presumably resulted in a more accurate characterisation of decision dynamics by reducing noise inherent in a simplistic, binary categorisation of feedback type. If further confirmed, this would further emphasise the need to incorporate neural measures into models of cognitive processing, as proposed by model-based cognitive neuroscience (Forstmann & Wagenmakers, 2015; Turner et al., 2015).

**Table S1.** Descriptive statistics for group-level DDM parameters. Mean, stand deviation, and the 2.5th and 97.5th quantiles for all group-level parameters in the best-fitting neurally-informed DDM. In this model, both the drift rate and boundary separation varied as a function of four EEG components (early positive, early negative, late positive, late negative) discriminant amplitudes from the previous trial.

| Parameter | $M$ | $SD$ | 2.5th & 97.5th<br>Quantiles |
| --- | --- | --- | --- |
| $a_0$ | 0.87 | 0.02 | [0.83, 0.92] |
| $a_{EarlyPos}$ | 0.02 | 0.006 | [0.003, 0.03] |
| $a_{EarlyNeg}$ | -0.007 | 0.007 | [-0.002, 0.006] |
| $a_{LatePos}$ | 0.01 | 0.005 | [0.0003, 0.02] |
| $a_{LateNeg}$ | -0.006 | 0.006 | [-0.002, 0.005] |
| $v_0$ | 0.70 | 0.07 | [0.57, 0.84] |
| $v_{EarlyPos}$ | 0.26 | 0.03 | [0.19, 0.33] |
| $v_{EarlyNeg}$ | 0.13 | 0.04 | [0.05, 0.21] |
| $v_{LatePos}$ | 0.29 | 0.03 | [0.22, 0.35] |
| $v_{LateNeg}$ | 0.09 | 0.04 | [0.02, 0.17] |
| $t$ | 0.43 | 0.009 | [0.41, 0.45] |
| *sv* | 0.62 | 0.17 | [0.23, 0.89] |
| *st* | 0.19 | 0.004 | [0.18, 0.20] |

## References

Akaike, H. (1978). A Bayesian analysis of the minimum AIC procedure. Annals of the Institute of Statistical Mathematics, 30(1), 9–14. doi:10.1007/BF02480194

Algermissen, J., Swart, J. C., Scheeringa, R., Cools, R., & den Ouden, H. E. M. (2024). Prefrontal signals precede striatal signals for biased credit assignment in motivational learning biases. Nature communications, 15(1). 10.1038/s41467-023-44632-x

Aston-Jones, G., & Cohen, J. D. (2005). An integrative theory of locus coeruleus-norepinephrine function: adaptive gain and optimal performance. Annu Rev Neurosci, 28, 403–450. doi:10.1146/annurev.neuro.28.061604.135709

Baayen, R. H., Davidson, D. J., & Bates, D. M. (2008). Mixed-effects modeling with crossed random effects for subjects and items. Journal of memory and language, 59(4), 390–412. doi:10.1016/j.jml.2007.12.005

Bland, A. R., & Schaefer, A. (2012). Different varieties of uncertainty in human decision-making. Front Neurosci, 6, 85. doi:10.3389/fnins.2012.00085

Botvinick, M. M., Braver, T. S., Barch, D. M., Carter, C. S., & Cohen, J. D. (2001). Conflict monitoring and cognitive control. Psychol Rev, 108(3), 624–652. doi:10.1037/0033-295x.108.3.624

Bouret, S., & Sara, S. J. (2005). Network reset: a simplified overarching theory of locus coeruleus noradrenaline function. Trends Neurosci, 28(11), 574–582. doi:10.1016/j.tins.2005.09.002

Briand, L. A., Gritton, H., Howe, W. M., Young, D. A., & Sarter, M. (2007). Modulators in concert for cognition: modulator interactions in the prefrontal cortex. Prog Neurobiol, 83(2), 69–91. doi:10.1016/j.pneurobio.2007.06.007

Brunello, N., Blier, P., Judd, L. L., Mendlewicz, J., Nelson, C. J., Souery, D.,…Racagni, G. (2003). Noradrenaline in mood and anxiety disorders: basic and clinical studies. Int Clin Psychopharmacol, 18(4), 191–202. doi:10.1097/00004850-200307000-00001

Carvalheiro, J., & Philiastides, M. G. (2023). Distinct spatiotemporal brainstem pathways of outcome valence during reward- and punishment-based learning. Cell Reports, 42(12). doi:ARTN 113589 10.1016/j.celrep.2023.113589

Cavanagh, J. F., Wiecki, T. V., Cohen, M. X., Figueroa, C. M., Samanta, J., Sherman, S. J., & Frank, M. J. (2011). Subthalamic nucleus stimulation reverses mediofrontal influence over decision threshold. Nat Neurosci, 14(11), 1462–1467. doi:10.1038/nn.2925

Chakroun, K., Mathar, D., Wiehler, A., Ganzer, F., & Peters, J. (2020). Dopaminergic modulation of the exploration/exploitation trade-off in human decision-making. Elife, 9. doi:10.7554/eLife.51260

Chakroun, K., Wiehler, A., Wagner, B., Mathar, D., Ganzer, F., van Eimeren, T.,…Peters, J. (2023). Dopamine regulates decision thresholds in human reinforcement learning in males. Nat Commun, 14(1), 5369. doi:10.1038/s41467-023-41130-y

Cinotti, F., Fresno, V., Aklil, N., Coutureau, E., Girard, B., Marchand, A. R., & Khamassi, M. (2019). Dopamine blockade impairs the exploration-exploitation trade-off in rats. Sci Rep, 9(1), 6770. doi:10.1038/s41598-019-43245-z

Colizoli, O., de Gee, J. W., Urai, A. E., & Donner, T. H. (2018). Task-evoked pupil responses reflect internal belief states. Sci Rep, 8(1), 13702. doi:10.1038/s41598-018-31985-3

Daw, N. D., O’Doherty, J. P., Dayan, P., Seymour, B., & Dolan, R. J. (2006). Cortical substrates for exploratory decisions in humans. Nature, 441(7095), 876–879. doi:10.1038/nature04766

Dayan, P., & Yu, A. J. (2006). Phasic norepinephrine: A neural interrupt signal for unexpected events. Network (Bristol), 17(4), 335–350. doi:10.1080/09548980601004024

de A Marcelino, A. L., Gray, O., Al-Fatly, B., Gilmour, W., Douglas Steele, J., Kuhn, A. A., & Gilbertson, T. (2023). Pallidal neuromodulation of the explore/exploit trade-off in decision-making. Elife, 12. doi:10.7554/eLife.79642

de Gee, J. W., Colizoli, O., Kloosterman, N. A., Knapen, T., Nieuwenhuis, S., & Donner, T. H. (2017). Dynamic modulation of decision biases by brainstem arousal systems. Elife, 6. doi:10.7554/eLife.23232

de Gee, J. W., Correa, C. M. C., Weaver, M., Donner, T. H., & van Gaal, S. (2021). Pupil Dilation and the Slow Wave ERP Reflect Surprise about Choice Outcome Resulting from Intrinsic Variability in Decision Confidence. Cereb Cortex, 31(7), 3565–3578. doi:10.1093/cercor/bhab032

de Gee, J. W., Knapen, T., & Donner, T. H. (2014). Decision-related pupil dilation reflects upcoming choice and individual bias. Proc Natl Acad Sci U S A, 111(5), E618–625. doi:10.1073/pnas.1317557111

Delorme, A., Sejnowski, T., & Makeig, S. (2007). Enhanced detection of artifacts in EEG data using higher-order statistics and independent component analysis. Neuroimage, 34(4), 1443–1449. doi:10.1016/j.neuroimage.2006.11.004

Denison, R. N., Parker, J. A., & Carrasco, M. (2020). Modeling pupil responses to rapid sequential events. Behav Res Methods, 52(5), 1991–2007. doi:10.3758/s13428-020-01368-6

Dutilh, G., Vandekerckhove, J., Forstmann, B. U., Keuleers, E., Brysbaert, M., & Wagenmakers, E. J. (2012). Testing theories of post-error slowing. Atten Percept Psychophys, 74(2), 454–465. doi:10.3758/s13414-011-0243-2

Ehlers, M. R., & Todd, R. M. (2017). Genesis and Maintenance of Attentional Biases: The Role of the Locus Coeruleus-Noradrenaline System. Neural Plast, 2017, 6817349. doi:10.1155/2017/6817349

Eldar, E., Cohen, J. D., & Niv, Y. (2013). The effects of neural gain on attention and learning. Nat Neurosci, 16(8), 1146–1153. doi:10.1038/nn.3428

Etkin, A., Egner, T., & Kalisch, R. (2011). Emotional processing in anterior cingulate and medial prefrontal cortex. Trends Cogn Sci, 15(2), 85–93. doi:10.1016/j.tics.2010.11.004

Filipowicz, A. L., Glaze, C. M., Kable, J. W., & Gold, J. I. (2020). Pupil diameter encodes the idiosyncratic, cognitive complexity of belief updating. Elife, 9. doi:10.7554/eLife.57872

Finke, J. B., Roesmann, K., Stalder, T., & Klucken, T. (2021). Pupil dilation as an index of Pavlovian conditioning. A systematic review and meta-analysis. Neurosci Biobehav Rev, 130, 351–368. doi:10.1016/j.neubiorev.2021.09.005

Fontanesi, L., Palminteri, S., & Lebreton, M. (2019). Decomposing the effects of context valence and feedback information on speed and accuracy during reinforcement learning: a meta-analytical approach using diffusion decision modeling. Cogn Affect Behav Neurosci, 19(3), 490–502. doi:10.3758/s13415-019-00723-1

Forstmann, B. U., Anwander, A., Schafer, A., Neumann, J., Brown, S., Wagenmakers, E. J.,…Turner, R. (2010). Cortico-striatal connections predict control over speed and accuracy in perceptual decision making. Proc Natl Acad Sci U S A, 107(36), 15916–15920. doi:10.1073/pnas.1004932107

Forstmann, B. U., Brass, M., Koch, I., & von Cramon, D. Y. (2006). Voluntary selection of task sets revealed by functional magnetic resonance imaging. J Cogn Neurosci, 18(3), 388–398. doi:10.1162/089892906775990589

Forstmann, B. U., Wagenmakers, E.-J., & SpringerLink. (2015). An introduction to model-based cognitive neuroscience. New York, NY: Springer.

Fouragnan, E., Queirazza, F., Retzler, C., Mullinger, K. J., & Philiastides, M. G. (2017). Spatiotemporal neural characterization of prediction error valence and surprise during reward learning in humans. Sci Rep, 7(1), 4762. doi:10.1038/s41598-017-04507-w

Fouragnan, E., Retzler, C., Mullinger, K., & Philiastides, M. G. (2015). Two spatiotemporally distinct value systems shape reward-based learning in the human brain. Nat Commun, 6, 8107. doi:10.1038/ncomms9107

Fouragnan, E., Retzler, C., & Philiastides, M. G. (2018). Separate neural representations of prediction error valence and surprise: Evidence from an fMRI meta-analysis. Hum Brain Mapp, 39(7), 2887–2906. doi:10.1002/hbm.24047

Fouragnan, E. F., Chau, B. K. H., Folloni, D., Kolling, N., Verhagen, L., Klein-Flügge, M.,…Rushworth, M. F. S. (2019). The macaque anterior cingulate cortex translates counterfactual choice value into actual behavioral change. Nature neuroscience, 22(5), 797-+. doi:10.1038/s41593-019-0375-6

Frank, M. J., Doll, B. B., Oas-Terpstra, J., & Moreno, F. (2009). Prefrontal and striatal dopaminergic genes predict individual differences in exploration and exploitation. Nat Neurosci, 12(8), 1062–1068. doi:10.1038/nn.2342

Frank, M. J., Gagne, C., Nyhus, E., Masters, S., Wiecki, T. V., Cavanagh, J. F., & Badre, D. (2015). fMRI and EEG predictors of dynamic decision parameters during human reinforcement learning. J Neurosci, 35(2), 485–494. doi:10.1523/JNEUROSCI.2036-14.2015

Franzen, L., Delis, I., De Sousa, G., Kayser, C., & Philiastides, M. G. (2020). Auditory information enhances post-sensory visual evidence during rapid multisensory decision-making. Nat Commun, 11(1), 5440. doi:10.1038/s41467-020-19306-7

Gelman, A., & Rubin, D. B. (1992). Inference from Iterative Simulation Using Multiple Sequences. Statistical science, 7(4), 457–472. doi:10.1214/ss/1177011136

Gerber E.M. (2025). permutest (https://www.mathworks.com/matlabcentral/fileexchange/71737-permutest), MATLAB Central File Exchange. Retrieved August 31, 2025.

Gilzenrat, M. S., Nieuwenhuis, S., Jepma, M., & Cohen, J. D. (2010). Pupil diameter tracks changes in control state predicted by the adaptive gain theory of locus coeruleus function. Cogn Affect Behav Neurosci, 10(2), 252–269. doi:10.3758/CABN.10.2.252

Glascher, J., Hampton, A. N., & O’Doherty, J. P. (2009). Determining a role for ventromedial prefrontal cortex in encoding action-based value signals during reward-related decision making. Cereb Cortex, 19(2), 483–495. doi:10.1093/cercor/bhn098

Goldfarb, S., Wong-Lin, K., Schwemmer, M., Leonard, N. E., & Holmes, P. (2012). Can post-error dynamics explain sequential reaction time patterns? Front Psychol, 3, 213. doi:10.3389/fpsyg.2012.00213

Hampton, A. N., Bossaerts, P., & O’Doherty, J. P. (2006). The role of the ventromedial prefrontal cortex in abstract state-based inference during decision making in humans. J Neurosci, 26(32), 8360–8367. doi:10.1523/JNEUROSCI.1010-06.2006

Harada, T. (2020). Learning From Success or Failure? - Positivity Biases Revisited. Frontiers in psychology, 11. 10.3389/fpsyg.2020.01627

Herrmann, N., Lanctot, K. L., & Khan, L. R. (2004). The role of norepinephrine in the behavioral and psychological symptoms of dementia. J Neuropsychiatry Clin Neurosci, 16(3), 261–276. doi:10.1176/jnp.16.3.261

Hoeks, B., & Levelt, W. J. M. (1993). Pupillary dilation as a measure of attention: A quantitative system analysis. Behavior research methods, instruments, & computers, 25(1), 16–26. doi:10.3758/BF03204445

Jepma, M., & Nieuwenhuis, S. (2011). Pupil diameter predicts changes in the exploration-exploitation trade-off: evidence for the adaptive gain theory. J Cogn Neurosci, 23(7), 1587–1596. doi:10.1162/jocn.2010.21548

Joshi, S., & Gold, J. I. (2020). Pupil Size as a Window on Neural Substrates of Cognition. Trends Cogn Sci, 24(6), 466–480. doi:10.1016/j.tics.2020.03.005

Joshi, S., Li, Y., Kalwani, R. M., & Gold, J. I. (2016). Relationships between Pupil Diameter and Neuronal Activity in the Locus Coeruleus, Colliculi, and Cingulate Cortex. Neuron, 89(1), 221–234. doi:10.1016/j.neuron.2015.11.028

Kluyver, Thomas, Ragan-Kelley, Benjamin, Pérez, Fernando, Granger, Brian, Bussonnier, Matthias, Frederic, Jonathan, Kelley, Kyle, Hamrick, Jessica, Grout, Jason, Corlay, Sylvain, Ivanov, Paul, Avila, Damián, Abdalla, Safia, Willing, Carol and Jupyter development team, (2016) Jupyter Notebooks – a publishing format for reproducible computational workflows. Loizides, Fernando and Scmidt, Birgit (eds.) In Positioning and Power in Academic Publishing: Players, Agents and Agendas. IOS Press. pp. 87–90. (doi:10.3233/978-1-61499-649-1-87).

Kolling, N., Behrens, T. E. J., Mars, R. B., & Rushworth, M. F. S. (2012). Neural Mechanisms of Foraging. Science, 336(6077), 95–98. doi:10.1126/science.1216930

Krajbich, I., Armel, C., & Rangel, A. (2010). Visual fixations and the computation and comparison of value in simple choice. Nat Neurosci, 13(10), 1292–1298. doi:10.1038/nn.2635

Krishnamurthy, K., Nassar, M. R., Sarode, S., & Gold, J. I. (2017). Arousal-related adjustments of perceptual biases optimize perception in dynamic environments. Nat Hum Behav, 1. doi:10.1038/s41562-017-0107

Krugel, L. K., Biele, G., Mohr, P. N., Li, S. C., & Heekeren, H. R. (2009). Genetic variation in dopaminergic neuromodulation influences the ability to rapidly and flexibly adapt decisions. Proc Natl Acad Sci U S A, 106(42), 17951–17956. doi:10.1073/pnas.090519110

Kruschke, J. K. (2011). Doing Bayesian data analysis: a tutorial with R and BUGS. Burlington, Mass: Academic Press.

Lavín, C., San Martin, R., & Rosales Jubal, E. (2014). Pupil dilation signals uncertainty and surprise in a learning gambling task. Front Behav Neurosci, 7, 218. doi:10.3389/fnbeh.2013.00218

Leonard, B. E. (1997). The role of noradrenaline in depression: a review. J Psychopharmacol, 11(4 Suppl), S39-47. Retrieved from https://www.ncbi.nlm.nih.gov/pubmed/9438232

Maness, E. B., Burk, J. A., McKenna, J. T., Schiffino, F. L., Strecker, R. E., & McCoy, J. G. (2022). Role of the locus coeruleus and basal forebrain in arousal and attention. Brain Res Bull, 188, 47–58. doi:10.1016/j.brainresbull.2022.07.014

Maris, E., & Oostenveld, R. (2007). Nonparametric statistical testing of EEG- and MEG-data. Journal of Neuroscience Methods, 164(1), 177–190. doi:10.1016/j.jneumeth.2007.03.024

Mathot, S., & Vilotijevic, A. (2022). Methods in cognitive pupillometry: Design, preprocessing, and statistical analysis. Behav Res Methods. doi:10.3758/s13428-022-01957-7

Mattes, A., Porth, E., & Stahl, J. (2022). Linking neurophysiological processes of action monitoring to post-response speed-accuracy adjustments in a neuro-cognitive diffusion model. Neuroimage, 247, 118798. doi:10.1016/j.neuroimage.2021.118798

Matzke, D., Dolan, C. V., Logan, G. D., Brown, S. D., & Wagenmakers, E. J. (2013). Bayesian parametric estimation of stop-signal reaction time distributions. J Exp Psychol Gen, 142(4), 1047–1073. doi:10.1037/a0030543

Matzke, D., & Wagenmakers, E. J. (2009). Psychological interpretation of the ex-Gaussian and shifted Wald parameters: a diffusion model analysis. Psychon Bull Rev, 16(5), 798–817. doi:10.3758/PBR.16.5.798

Murphy, K., & Fouragnan, E. (2024). The future of transcranial ultrasound as a precision brain interface. Plos Biology, 22(10). doi:ARTN e3002884 10.1371/journal.pbio.3002884

Murphy, P. R., O’Connell, R. G., O’Sullivan, M., Robertson, I. H., & Balsters, J. H. (2014). Pupil diameter covaries with BOLD activity in human locus coeruleus. Hum Brain Mapp, 35(8), 4140–4154. doi:10.1002/hbm.22466

Murphy, P. R., van Moort, M. L., & Nieuwenhuis, S. (2016). The Pupillary Orienting Response Predicts Adaptive Behavioral Adjustment after Errors. PLoS One, 11(3), e0151763. doi:10.1371/journal.pone.0151763

Nassar, M. R., Rumsey, K. M., Wilson, R. C., Parikh, K., Heasly, B., & Gold, J. I. (2012). Rational regulation of learning dynamics by pupil-linked arousal systems. Nat Neurosci, 15(7), 1040–1046. doi:10.1038/nn.3130

Navarro, D. J., Fuss, I.G. (2009). Fast and accurate calculations for first-passage times in Wiener diffusion models. Journal of mathematical psychology, 53(4), 222–230. 10.1016/j.jmp.2009.02.003

Nunez, M. D., Vandekerckhove, J., & Srinivasan, R. (2017). How attention influences perceptual decision making: Single-trial EEG correlates of drift-diffusion model parameters. Journal of mathematical psychology, 76(Pt B), 117–130. doi:10.1016/j.jmp.2016.03.003

O’Doherty, J. P., Hampton, A., & Kim, H. (2007). Model-based fMRI and its application to reward learning and decision making. Ann N Y Acad Sci, 1104, 35–53. doi:10.1196/annals.1390.022

Pajkossy, P., Szollosi, A., Demeter, G., & Racsmany, M. (2017). Tonic noradrenergic activity modulates explorative behavior and attentional set shifting: Evidence from pupillometry and gaze pattern analysis. Psychophysiology, 54(12), 1839–1854. doi:10.1111/psyp.12964

Palminteri, S., Wyart, V., & Koechlin, E. (2017). The Importance of Falsification in Computational Cognitive Modeling. Trends in Cognitive Sciences, 21(6), 425–433. doi:10.1016/j.tics.2017.03.011

Parra, L. C., Spence, C. D., Gerson, A. D., & Sajda, P. (2005). Recipes for the linear analysis of EEG. Neuroimage, 28(2), 326–341. doi:10.1016/j.neuroimage.2005.05.032

Pedersen, M. L., & Frank, M. J. (2020). Simultaneous Hierarchical Bayesian Parameter Estimation for Reinforcement Learning and Drift Diffusion Models: a Tutorial and Links to Neural Data. Comput Brain Behav, 3(4), 458–471. doi:10.1007/s42113-020-00084-w

Pedersen, M. L., Frank, M. J., & Biele, G. (2017). The drift diffusion model as the choice rule in reinforcement learning. Psychonomic Bulletin & Review, 24(4), 1234–1251. doi:10.3758/s13423-016-1199-y

Pernet, C. R., Wilcox, R., & Rousselet, G. A. (2012). Robust correlation analyses: false positive and power validation using a new open source matlab toolbox. Front Psychol, 3, 606. doi:10.3389/fpsyg.2012.00606

Philiastides, M. G., Ratcliff, R., & Sajda, P. (2006). Neural representation of task difficulty and decision making during perceptual categorization: a timing diagram. J Neurosci, 26(35), 8965–8975. doi:10.1523/JNEUROSCI.1655-06.2006

Philiastides, M. G., & Sajda, P. (2006). Temporal characterization of the neural correlates of perceptual decision making in the human brain. Cereb Cortex, 16(4), 509–518. doi:10.1093/cercor/bhi130

Preuschoff, K., t Hart, B. M., & Einhauser, W. (2011). Pupil Dilation Signals Surprise: Evidence for Noradrenaline’s Role in Decision Making. Front Neurosci, 5, 115. doi:10.3389/fnins.2011.00115

Filippo Queirazza, Marios Philiastides. Prefrontal and parieto-occipital neural signatures of evidence accumulation and response to computerised Cognitive Behavioural Therapy in depression., 19 December 2024, PREPRINT (Version 1) available at Research Square [10.21203/rs.3.rs-5615710/v1]

Ratcliff, R. (1978). A theory of memory retrieval. Psychological Review, 85(2), 59–108. doi:10.1037/0033-295X.85.2.59

Ratcliff, R., & McKoon, G. (2008). The diffusion decision model: theory and data for two-choice decision tasks. Neural Comput, 20(4), 873–922. doi:10.1162/neco.2008.12-06-420

Ratcliff, R., Philiastides, M. G., & Sajda, P. (2009). Quality of evidence for perceptual decision making is indexed by trial-to-trial variability of the EEG. Proc Natl Acad Sci U S A, 106(16), 6539–6544. doi:10.1073/pnas.0812589106

Ratcliff, R., & Rouder, J. N. (1998). Modeling Response Times for Two-Choice Decisions. Psychological science, 9(5), 347–356. doi:10.1111/1467-9280.00067

Reimer, J., McGinley, M. J., Liu, Y., Rodenkirch, C., Wang, Q., McCormick, D. A., & Tolias, A. S. (2016). Pupil fluctuations track rapid changes in adrenergic and cholinergic activity in cortex. Nat Commun, 7, 13289. doi:10.1038/ncomms13289

Rushworth, M. F., & Behrens, T. E. (2008). Choice, uncertainty and value in prefrontal and cingulate cortex. Nat Neurosci, 11(4), 389–397. doi:10.1038/nn2066

Sajda, P., Philiastides, M. G., & Parra, L. C. (2009). Single-trial analysis of neuroimaging data: inferring neural networks underlying perceptual decision-making in the human brain. IEEE Rev Biomed Eng, 2, 97–109. doi:10.1109/RBME.2009.2034535

Sara, S. J. (2009). The locus coeruleus and noradrenergic modulation of cognition. Nat Rev Neurosci, 10(3), 211–223. doi:10.1038/nrn2573

Sara, S. J., & Bouret, S. (2012). Orienting and reorienting: the locus coeruleus mediates cognition through arousal. Neuron, 76(1), 130–141. doi:10.1016/j.neuron.2012.09.011

Schultz, W. (2016). Dopamine reward prediction-error signalling: a two-component response. Nature Reviews Neuroscience, 17(3), 183–195. doi:10.1038/nrn.2015.26

Schultz, W., Dayan, P., & Montague, P. R. (1997). A neural substrate of prediction and reward. Science, 275(5306), 1593–1599. doi:10.1126/science.275.5306.1593

Schwarz, G. (1978). Estimating the Dimension of a Model. The Annals of statistics, 6(2), 461–464. doi:10.1214/aos/1176344136

The Mathworks Inc. (2018). MATLAB version 9.5.0.944444 (R2018b). Natick, Massachusetts: The Mathworks Inc. https://www.mathworks.com

Turner, B. M., van Maanen, L., & Forstmann, B. U. (2015). Informing cognitive abstractions through neuroimaging: the neural drift diffusion model. Psychol Rev, 122(2), 312–336. doi:10.1037/a0038894

Spiegelhalter, D. J., Best, N. G., Carlin, B. R., & van der Linde, A. (2002). Bayesian measures of model complexity and fit. Journal of the Royal Statistical Society Series B-Statistical Methodology, 64, 583–616. doi:Doi 10.1111/1467-9868.00353

Torregrossa, M. (2019). Neural mechanisms of addiction. London: Academic Press.

Ullsperger, M., & von Cramon, D. Y. (2001). Subprocesses of performance monitoring: a dissociation of error processing and response competition revealed by event-related fMRI and ERPs. Neuroimage, 14(6), 1387–1401. doi:10.1006/nimg.2001.0935

Urai, A. E., Braun, A., & Donner, T. H. (2017). Pupil-linked arousal is driven by decision uncertainty and alters serial choice bias. Nat Commun, 8, 14637. doi:10.1038/ncomms14637

van Rij, J., Hendriks, P., van Rijn, H., Baayen, R. H., & Wood, S. N. (2019). Analyzing the Time Course of Pupillometric Data. Trends Hear, 23, 2331216519832483. doi:10.1177/2331216519832483

van Rossum, G., & Drake, F. L. (2009). Python 3 Reference Manual. Scotts Valley, CA: CreateSpace.

Van Slooten, J. C., Jahfari, S., Knapen, T., & Theeuwes, J. (2017). Individual differences in eye blink rate predict both transient and tonic pupil responses during reversal learning. PLoS One, 12(9), e0185665. doi:10.1371/journal.pone.0185665

Van Slooten, J. C., Jahfari, S., Knapen, T., & Theeuwes, J. (2018). How pupil responses track value-based decision-making during and after reinforcement learning. PLoS Comput Biol, 14(11), e1006632. doi:10.1371/journal.pcbi.1006632

Van Slooten, J. C., Jahfari, S., & Theeuwes, J. (2019). Spontaneous eye blink rate predicts individual differences in exploration and exploitation during reinforcement learning. Sci Rep, 9(1), 17436. doi:10.1038/s41598-019-53805-y

Varazzani, C., San-Galli, A., Gilardeau, S., & Bouret, S. (2015). Noradrenaline and dopamine neurons in the reward/effort trade-off: a direct electrophysiological comparison in behaving monkeys. J Neurosci, 35(20), 7866–7877. doi:10.1523/JNEUROSCI.0454-15.2015

Warren, C. M., Wilson, R. C., van der Wee, N. J., Giltay, E. J., van Noorden, M. S., Cohen, J. D., & Nieuwenhuis, S. (2017). The effect of atomoxetine on random and directed exploration in humans. PLoS One, 12(4), e0176034. doi:10.1371/journal.pone.0176034

Weiss, A. R., Gillies, M. J., Philiastides, M. G., Apps, M. A., Whittington, M. A., FitzGerald, J. J.,…Green, A. L. (2018). Dorsal Anterior Cingulate Cortices Differentially Lateralize Prediction Errors and Outcome Valence in a Decision-Making Task. Frontiers in human neuroscience, 12. doi:ARTN 203 10.3389/fnhum.2018.00203

Westwood, S., & Philiastides, M. G. (2024). Early Salience Signals Predict Interindividual Asymmetry in Decision Accuracy Across Rewarding and Punishing Contexts. Human Brain Mapping, 45(17). doi:ARTN e70072 10.1002/hbm.70072

Wiecki, T. V., Sofer, I., & Frank, M. J. (2013). HDDM: Hierarchical Bayesian estimation of the Drift-Diffusion Model in Python. Front Neuroinform, 7, 14. doi:10.3389/fninf.2013.00014

Yu, A. J., & Dayan, P. (2005). Uncertainty, neuromodulation, and attention. Neuron, 46(4), 681–692. doi:10.1016/j.neuron.2005.04.026

Zenon, A. (2019). Eye pupil signals information gain. Proc Biol Sci, 286(1911), 20191593. doi:10.1098/rspb.2019.1593

Zhang, J., & Rowe, J. B. (2014). Dissociable mechanisms of speed-accuracy tradeoff during visual perceptual learning are revealed by a hierarchical drift-diffusion model. Front Neurosci, 8, 69. doi:10.3389/fnins.2014.00069

